# Tumor cell-intrinsic mtDNA instability drives cGAS-dependent temporal remodeling of the melanoma immune microenvironment

**DOI:** 10.64898/2026.09.28.755003

**Authors:** Abate Assefa Bashaw, Ashley D. Winters, Yuanjiu Lei, Luiza Ferreira Araujo, Do Gia Tam Vo, Arunachalam Sekar, John F. Bachman, Paige S. Graves, Melissa Pineda, Jill C. Rubinstein, Gerald S. Shadel, Marcus Bosenberg, Laura Ciaccia West, A. Phillip West

**Author notes:** Correspondence: A. Phillip West, PhD, The Jackson Laboratory, 600 Main Street, Bar Harbor, ME, 06409, USA.

## Abstract

Mitochondrial DNA (mtDNA) instability is an emerging feature of cancer, yet its biological effects on the tumor microenvironment (TME) and tumor progression remain incompletely defined. Using melanocyte-specific conditional silencing of mitochondrial transcription factor A (TFAM) in an autochthonous *Braf^V600E^*/*Pten^-/-^* melanoma model, we demonstrate that tumor cell-intrinsic mtDNA instability accelerates melanoma growth and metastatic dissemination in a cGAS-dependent manner. Transcriptomic profiling of early and late tumor stages revealed that mtDNA instability drives temporal remodeling of the tumor microenvironment (TME), characterized by an early type I interferon (IFN)-rich inflammatory state that licenses immunoregulatory programs as tumors progress. Tumor-associated macrophages emerged as the principal immune cell type reshaped by tumor cell-intrinsic mtDNA stress signaling, with early enrichment of inflammatory states and a shift toward efferocytosis-associated, tissue-adaptive states. The early mtDNA instability-associated macrophage signatures mapped to human melanoma states associated with favorable survival and therapy response, whereas the late signatures mapped to states associated with poor prognosis and therapy resistance across independent cohorts. Anti-PD-1 treatment delayed progression of mtDNA-instable tumors, consistent with a therapeutic sensitivity window associated with the early IFN-rich immune state. These findings establish tumor cell-intrinsic mtDNA instability as a driver of temporal immune-stromal remodeling that promotes melanoma progression and shapes therapeutic response.

## Introduction

Mitochondrial DNA (mtDNA) instability, including somatic mutations and copy number alterations, is a common molecular feature of cancer and is associated with poor clinical outcomes across tumor types ^1,2^. mtDNA stress can arise through oxidative damage, replication errors, or disruption of nuclear-encoded regulators of mtDNA maintenance, including mitochondrial transcription factor A (TFAM), the mtDNA polymerase POLG, and the mtDNA helicase Twinkle ^3–5^.

Melanoma arises from melanocytes chronically exposed to ultraviolet radiation, which induces oxidative mtDNA damage that can drive mtDNA instability ^6,7^, features frequently detected in advanced and metastatic melanoma ^8^ and increasingly observed with aging across human cancers ^2^. Although mtDNA perturbations have been linked to tumorigenesis, their biological consequences remain highly context-dependent and often conflicting. Melanoma cell-derived mutant mtDNA transferred to tumor-infiltrating T cells impairs mitochondrial fitness and reduces response to immune checkpoint blockade (ICB) ^9^, while patients carrying germline mitochondrial haplogroup T variants similarly exhibit reduced ICB responses in metastatic melanoma ^10^. Conversely, truncating mutations in *mt-Nd5-encoded* complex I subunit that impairs oxidative phosphorylation (OXPHOS) enhances ICB sensitivity ^11^, whereas pathogenic mtDNA variants introduced into melanoma cybrid lines reduce metastatic dissemination in immunodeficient mice ^12^. Under mitochondrial stress, mitochondria release damage-associated molecular patterns (DAMPs), including mtDNA, which can activate innate immune pathways, most notably the cyclic GMP-AMP synthase-stimulator of interferon (IFN) genes (cGAS-STING) pathway, triggering inflammatory responses ^3^. Although generally considered antitumoral, cGAS-STING signaling has been shown to exert pro-tumorigenic effects ^13,14^. Because mtDNA is present in multiple copies per cell, individual tumor cells can harbor mtDNA variants at different levels of heteroplasmy, resulting in variable degrees of mtDNA stress and functional consequences among tumor cells ^15^. Defining the causal contribution of mtDNA instability to tumor progression and antitumor immunity therefore is complex and remains incompletely understood.

Using an autochthonous Braf^V600E^/Pten^-/-^ melanoma model with melanocyte-specific conditional TFAM knockdown, we investigated the effect of tumor cell-intrinsic mtDNA instability during melanoma progression. TFAM is the master regulator of mtDNA packaging and maintenance, and heterozygous ablation induces mtDNA instability without overt respiratory collapse ^3^. Reduced TFAM expression has also been associated with tumor progression and metastatic potential in human cancers, including melanoma ^16,17^. In this study, we demonstrated that tumor cell-intrinsic mtDNA instability promotes melanoma progression through cGAS-dependent innate immune activation. mtDNA instability drives temporal remodeling of the tumor immune microenvironment, with an early type I IFN-dominated response giving way to a late immunoregulatory state. Macrophage transcriptional signatures associated with this mtDNA instability-driven temporal remodeling stratified survival and therapeutic responses in human melanoma cohorts. Together, these findings establish that tumor cell-intrinsic mtDNA instability drives immune-stromal rewiring that promotes melanoma progression.

## Results

### TFAM silencing in melanoma cells *in vivo* promotes tumor progression and metastasis

Melanocyte-specific Tfam haploinsufficiency was established in an autochthonous melanoma mouse model driven by Braf^V600E^ activation and Pten loss using a CreER inducible system under the melanocyte-specific tyrosinase (*Tyr*) promoter ^18^. Mice carrying Braf^CA/+^, Pten^fl/fl^, and Tyr::CreER alleles were crossed with Tfam^fl/+^ mice to generate melanocyte-specific Tfam heterozygous deletion (TFAM^KD-MEL^) or Tfam wild-type (TFAM^WT-MEL^) control mice (Fig. 1a). Melanoma was induced by topical administration of 4-hydroxytamoxifen (4-OHT), leading to melanocyte-specific Cre recombination, Braf^V600E^ activation and Pten deletion, with concurrent heterozygous Tfam deletion in TFAM^KD-MEL^ mice (Fig. 1a; see Methods). PCR-based genotyping and immunoblotting of early-stage tumors confirmed Cre-mediated Tfam recombination (Extended Data Fig. 1a) and reduced TFAM protein levels in TFAM^KD-MEL^ tumors relative to TFAM^WT-MEL^ controls, despite the presence of Tfam-sufficient stromal and immune cells (Extended Data Fig. 1b). TFAM^KD-MEL^ tumors also exhibited reduced mtDNA copy number (Fig. 1b), confirming tumor cell–intrinsic mtDNA instability in vivo. Tumors were induced in young animals, enabling isolation of the consequences of tumor cell–intrinsic mtDNA instability from age-associated changes, including chromosomal instability. While tumor initiation kinetics were comparable between genotypes, TFAM^KD-MEL^ mice exhibited accelerated tumor growth and greater metastatic burden relative to TFAM^WT-MEL^ controls (Fig. 1c-e). TFAM^KD-MEL^ mice developed a higher incidence of secondary cutaneous lesions at both proximal and anatomically distal sites, including the tail and upper dorsal skin (Fig. 1c, d), consistent with increased multifocal cutaneous dissemination recapitulating satellite and in-transit metastases observed in high-risk stage III human melanoma ^19^. TFAM^KD-MEL^ tumors exhibited a shorter doubling time and accelerated growth (Fig. 1e; Extended Data Fig. 1c, d), and survival analysis from the time tumors became palpable to the humane endpoint revealed reduced overall and metastasis-free survival in TFAM^KD-MEL^ mice relative to controls (Fig. 1f, g). Histological examination revealed no significant differences in tumor morphology between genotypes (Extended Data Fig. 1e). Histological analysis of tumor-draining sentinel lymph nodes revealed increased numbers of pigmented cells in TFAM^KD-MEL^ mice relative to TFAM^WT-MEL^ controls (Fig. 1h, i), and immunofluorescence staining for the pan-melanoma marker SOX10 confirmed increased melanoma cell infiltration, distinguishing tumor cells from pigment-laden macrophages (melanophages) (Fig.1h, i). Together, these findings demonstrate that tumor cell–intrinsic Tfam ablation drives mtDNA instability–associated tumor growth and metastatic dissemination without affecting melanoma initiation.

**Fig. 1:**
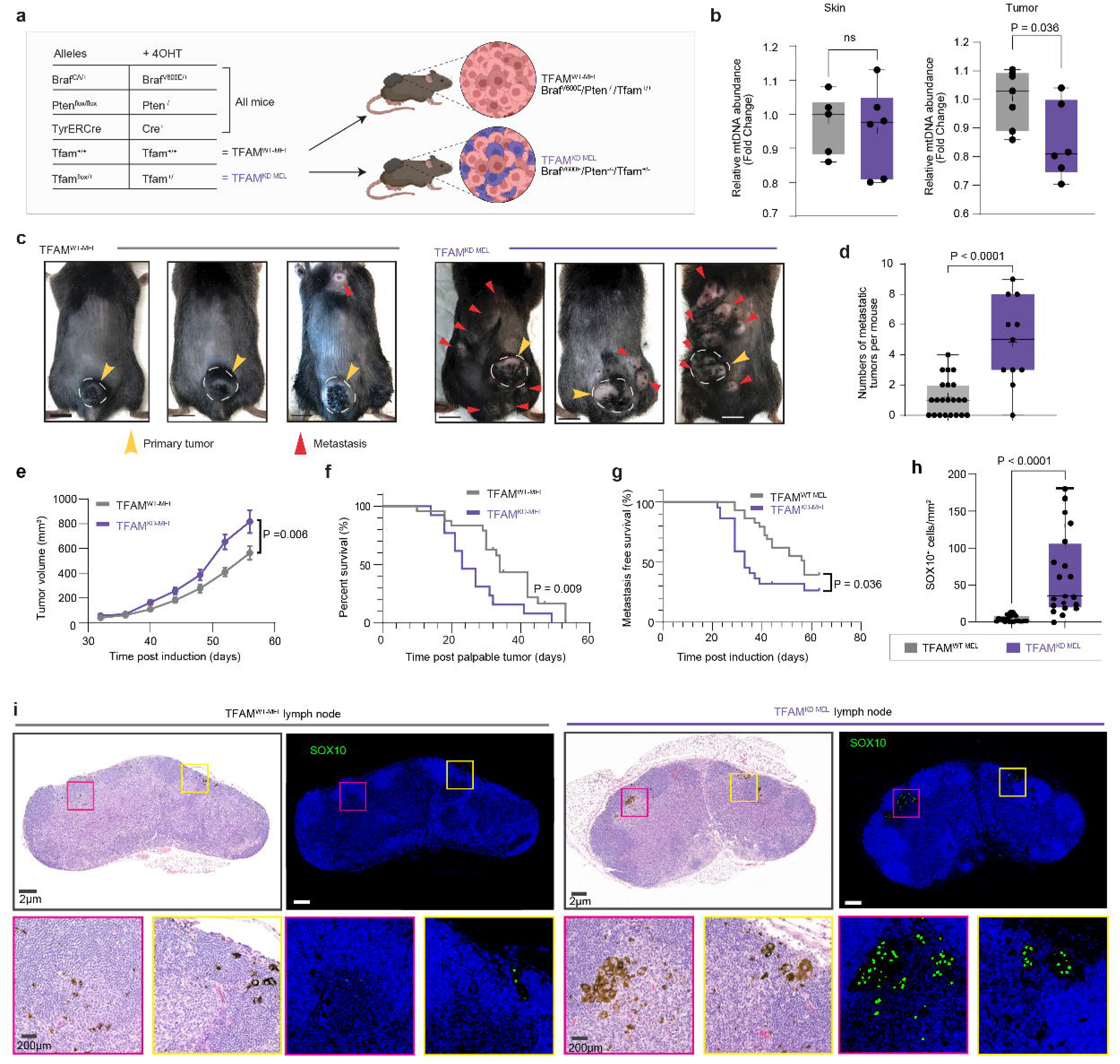
TFAM silencing-induced mtDNA instability accelerates melanoma growth and metastasis. a, Schematic of the Tfam knockdown melanoma model. Mice carrying the Braf^CA^, Pten^fl/fl^, and TyrERCre alleles were crossed with Tfam^fl/+^ mice and induced with 4-hydroxytamoxifen (4-OHT) to generate melanocyte-specific Tfam knockdown (TFAM^KD-MEL^) or Tfam wildtype (TFAM^WT-MEL^) tumors on a Braf^V600E^/Pten^-/-^ melanoma background. b, Relative mitochondrial DNA (mtDNA) abundance measured by mitochondrial gene copy number normalized to the nuclear reference gene Tert in skin prior to tumor induction (left) and established tumors (right). c, Representative images of tumor-bearing mice (n = 50-60 per genotype). Yellow arrowheads indicate primary tumors, and red arrowheads indicate metastatic lesions. d, Quantification of macroscopic metastatic lesions per mouse at experimental endpoint. e, Tumor growth kinetics of TFAM^WT-MEL^ and TFAM^KD-MEL^ mice (n = 50-60 per genotype). f,g, Kaplan-Meier survival analysis showing overall survival following palpable tumor detection (f) and metastasis-free survival (g). Statistical significance was determined using the log-rank (Mantel-Cox) test. h, Quantification of SOX10 cells per mm² in tumor-draining inguinal lymph nodes (iLNs) (n = 25-30 per genotype). Statistical significance was determined using two-way ANOVA with multiple comparisons correction. i, Representative hematoxylin and eosin (H&E) staining (left) and SOX10 immunofluorescence staining (green) with DAPI nuclear counterstaining (blue) (right) of tumor-draining iLNs. Scale bars, 200 μm.

### Melanoma cell-intrinsic mtDNA instability drives innate immune activation and inflammatory remodeling of the TME

To define the cell-intrinsic consequences of mtDNA instability in melanoma cells, we generated Tfam knockdown in YUMM1.7 melanoma cells using CRISPR-Cas9 with two independent single guide RNAs targeting *Tfam* (sgTFAM YUMM1.7) and a non-targeting control sgRNA (sgCTL YUMM1.7). YUMM1.7 is a syngeneic murine melanoma cell line derived from Braf^V600E^/Pten^-/-^ mice with additional Cdkn2a deletion. TFAM downregulation in sgTFAM YUMM1.7 cells was confirmed by immunofluorescence and immunoblotting (Extended Data Fig. 2a, b). TFAM knockdown induced mtDNA nucleoid aggregation and reduced mtDNA copy number (Extended Data Fig. 2a, d) without significant changes in core OXPHOS complex subunits (Extended Data Fig. 2c), supporting the use of sgTFAM YUMM1.7 cells as an in vitro model of mtDNA stress. Consistent with activation of the cGAS-STING pathway, sgTFAM YUMM1.7 cells showed increased levels of cGAS, STING1 and IRF3, together with phosphorylated TBK1 (Extended Data Fig. 2b). RNA sequencing (RNA-seq) of early passage sgTFAM YUMM1.7 cells further revealed robust induction of type I IFN signaling and innate immune programs (Fig. 2a, b). Upregulated genes included canonical IFN-stimulated genes (ISGs), such as *Isg15*, *Mx1*, *Ddx58*, *Ifih1*, and components of the major histocompatibility complex class I (MHC-I) antigen-presentation machinery (Fig. 2a), consistent with a cell-intrinsic mtDNA instability-driven type I IFN response. Pathway analysis confirmed enrichment of IFN signaling and MHC-I antigen presentation in sgTFAM YUMM1.7 cells (Fig. 2b). In parallel, sgTFAM YUMM1.7 cells upregulated genes associated with immune modulation (including *Cd274*, *Lgals9*, and *Sirpa*), as well as invasion and cellular plasticity including *Aldh1a3*, *Mmp2*, and WNT pathway components (Fig. 2a, b), suggesting activation of stress-adaptive programs that may support tumor cell survival and invasive capacity under persistent IFN signaling.

**Fig. 2:**
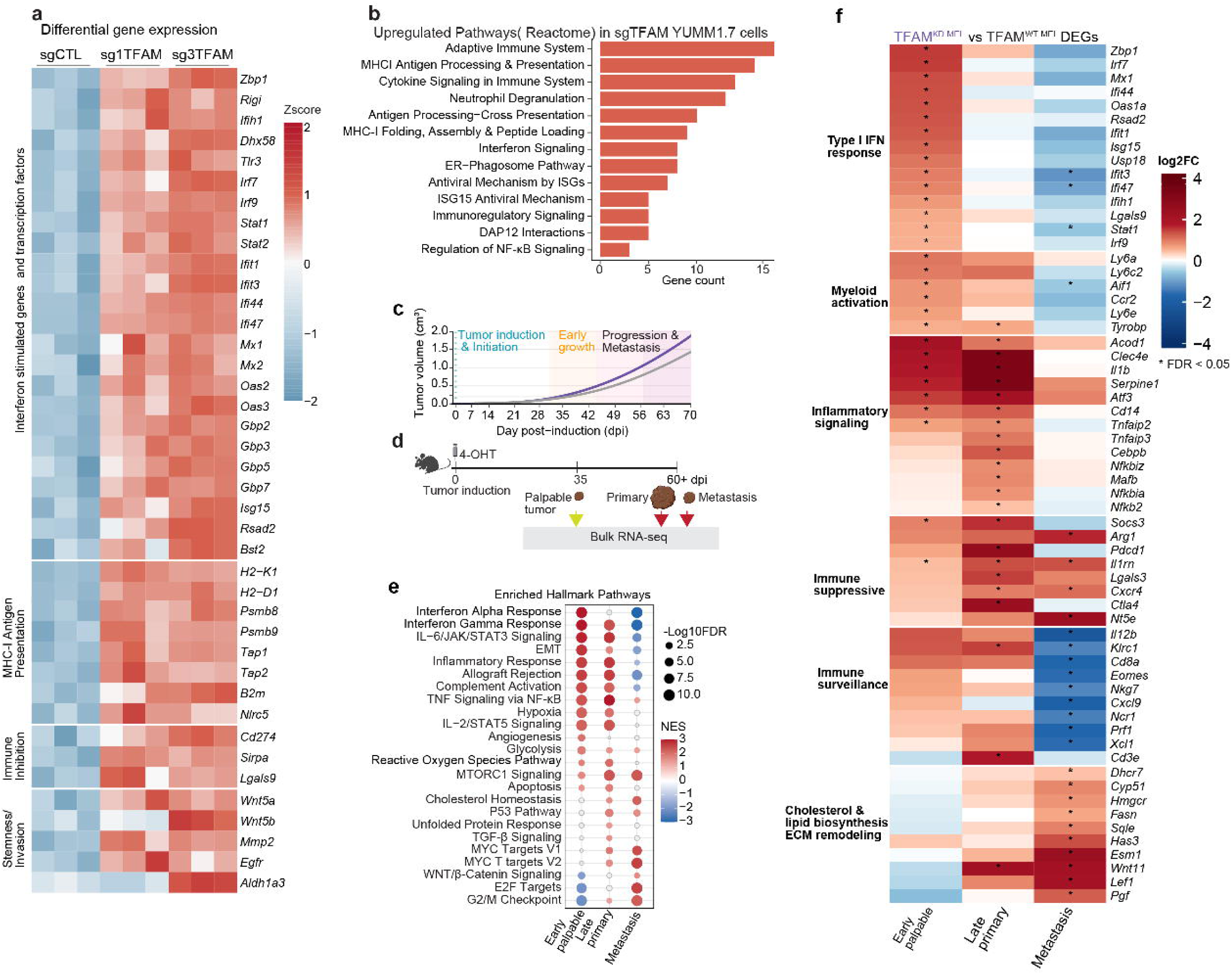
**Tumor-intrinsic mtDNA instability drives innate immune activation that fosters an immunosuppressive inflammatory TME**. a, Heatmap of differentially expressed genes (DEGs) in sg1TFAM and sg3TFAM YUMM1.7 cells relative to sgCTL control cells; values represent row-normalized z-scores. Genes with fold change > 1.2 and adjusted *P* < 0.05 are shown. b, Reactome pathways significantly enriched among DEGs shared between sg1TFAM and sg3TFAM cells relative to sgCTL controls. c,d, Schematic of the in vivo melanoma profiling strategy. c, Tumor growth kinetics. d, Experimental timeline. Melanoma was induced by topical application of 4-hydroxytamoxifen (4-OHT) on day 0, and bulk RNA sequencing was performed on early palpable tumors (35 days post-induction (dpi)), late primary tumors (1,000–1,500 mm³), and metastatic skin tumors (56–60 dpi). e, Hallmark pathway enrichment analysis of DEGs from early palpable, late primary, and metastatic TFAM^KD-MEL^ tumors relative to TFAM^WT-MEL^ tumors (FDR < 0.05). f, Joint heatmap showing log fold changes of DEGs in early palpable, late primary, and metastatic TFAM^KD-MEL^ tumors relative to TFAM^WT-MEL^ tumors (FDR < 0.05).

Beyond tumor cell-intrinsic effects, mtDNA stress can engage in autocrine and paracrine signaling that reshapes the TME. We therefore performed bulk RNA-seq of early palpable pigmented lesions and, at experimental endpoint, primary and metastatic tumors from TFAM^WT-MEL^ and TFAM^KD-MEL^ mice (Fig. 2c, d). This analysis revealed stage-dependent immune remodeling in TFAM^KD-MEL^ tumors, characterized by an early type I IFN response and a late inflammatory transcriptional state (Fig. 2e, f; Extended Data Fig. 2e-h).

Early palpable TFAM^KD-MEL^ lesions displayed robust type I IFN pathway activation that closely mirrored the transcriptional response of sgTFAM YUMM1.7 cells, including induction of canonical ISGs (e.g., *Zbp1, Irf7, Isg15, Mx1, Ifit1,* and *Ifit3*) and IFN-inducible negative regulators (*Socs3, Usp18,* and *Dhx58*) (Fig. 2e; Extended Data Fig. 2e, g). Pathway enrichment further confirmed activation of cytosolic DNA-sensing and IFN-α response programs (Fig. 2e; Extended Data Fig. 2h). Increased protein expression of cGAS, ZBP1, and IFIT3 provided independent evidence of engagement of the cGAS-STING-IFN axis in early TFAM^KD-MEL^ tumors (Extended Data Fig. 2f). Genes associated with myeloid cell recruitment, ECM remodeling and fibrosis, including *Csf1, Lgals3bp, Postn, Adamts2,* and *Mmp11*, were upregulated in both early TFAM^KD-MEL^ lesions and sgTFAM YUMM1.7 cells (Extended Data Fig. 2e), identifying tumor-associated programs that may contribute to subsequent immune and stromal remodeling. Consistent with this, early TFAM^KD-MEL^ tumors showed enrichment of complement activation and phagocytic programs, together with increased expression of myeloid recruitment and activation genes including *Ccr2, Aif1, Ly6c2,* and *Tyrobp*, as well as IFN-responsive myeloid genes *Ly6a* and *Ly6e* (Extended Data Fig. 2e, g). Upregulation of immunoregulatory genes such as Lilrb4a (Extended Data Fig. 2g) further indicated engagement of counter-regulatory programs alongside the early type I IFN response.

At late stage, the mtDNA instability-associated IFN response was largely replaced by a chronic inflammatory state. Compared with TFAM^WT-MEL^ tumors, late-stage TFAM^KD-MEL^ tumors showed increased expression of inflammatory mediators, including *Il1b*, *Atf3*, *Acod1* and *Serpine1*, and NF-κB-associated genes such as *Tnfaip3*, *Nfkbiz*, *Nfkbia* and *Nfkb2* (Fig. 2f; Extended Data Fig. 2i). TNF/NF-κB, IL-6/JAK/STAT3, and TGF-β signaling, as well as reactive oxygen species programs, were enriched in TFAM^KD-MEL^ tumors (Fig. 2e), indicating a shift toward NF-κB- and JAK/STAT3 -associated inflammatory signaling. Increased expression of immunoregulatory genes, including *Arg1*, *Pdcd1*, *Nt5e* and *Lgals3*, further supported an immunoregulatory TME. These findings are consistent with evidence of chronic type I IFN signaling and rewiring toward NF-κB- and JAK/STAT3-driven inflammatory programs that promote an immunoregulatory and pro-metastatic TME in the context of chromosomal instability (Hong et al., 2022; Li et al., 2023). Metastatic TFAM^KD-MEL^ tumors exhibited a further shift toward immune attenuation coupled with tumor cell-intrinsic invasive and proliferative programs (Fig. 2e, f). IFN-α and IFN-γ responses and complement activation, prominent in early palpable TFAM^KD-MEL^ lesions, were downregulated in TFAM^KD-MEL^ metastatic lesions (Fig. 2e). Consistent with late primary tumors, TFAM^KD-MEL^ metastatic lesions showed elevated expression of immunoregulatory genes including *Arg1*, *Pdcd1*, *Nt5e*, and *Lgals3*, while antitumor immune effector genes including *Cd8a*, *Prf1*, *Nkg7*, *Eomes*, and *Cxcl9* were among the most strongly downregulated transcripts relative to TFAM^WT-MEL^ metastatic lesions (Fig. 2e, f, Extended Data Fig. 2i, j), indicating an immune-excluded metastatic lesion. TFAM^KD-MEL^ metastatic tumors also exhibited enrichment of programs associated with tumor growth and invasion, including cholesterol and lipid biosynthesis, WNT signaling, and MYC targets, E2F targets, and G2/M checkpoint programs (Fig. 2e, f). These data indicate that mtDNA instability accompanies the acquisition of proliferative, invasive, and immune-evasive programs in metastatic lesions, consistent with reports that sustained type I IFN signaling can induce tumor-cell-intrinsic adaptations that attenuate cytotoxic immune responses (Musella et al., 2022; Qiu et al., 2023) Altogether the bulk transcriptomics suggest that mtDNA instability drives inflammatory TME that promotes tumor growth and metastasis.

### Single-cell transcriptomic profiling reveals that mtDNA instability in tumor cells drives temporal TME reprogramming

Bulk transcriptomic profiling established that mtDNA instability drives innate immune activation and inflammatory remodeling of TME in TFAM^KD-MEL^ tumors. To define the cellular basis of these changes, we performed single-cell RNA-seq (scRNA-seq) on TFAM^KD-MEL^ and TFAM^WT-MEL^ melanoma tumors at early (∼300-500 mm³, 42 days post-induction) and late (∼1,000-1,500 mm³, 55-60 days post-induction) stages (Fig. 3a). Seurat based integration and clustering identified malignant, myeloid, lymphoid, and stromal compartments across 23 distinct clusters, which were annotated into major cell types based on canonical marker expression (Fig. 3b, c; Extended Data Fig.3a, b). Early TFAM^KD-MEL^ TME exhibited marked immune remodeling, with increased macrophage abundance (37.4% versus 18.0%) and reduced regulatory T cells (29.7% versus 40.9%) relative to TFAM^WT-MEL^ controls (Fig. 3c). B cells were also expanded in early TFAM^KD-MEL^ tumors, predominantly reflecting pre- and pro-B progenitor cell states (Fig. 3c; Extended Data Fig. 6a, b). At late stage, TFAM^KD-MEL^ tumors showed a marked increase in an immature myeloid cell population (IMC; 42.3% versus 16.6%) (Fig. 3c; Extended Data Fig. 6), indicating myeloid cell remodeling, promoting suppressive TME. Hallmark pathway gene set enrichment analysis across major cell types revealed broad induction of inflammatory signaling in early TFAM^KD-MEL^ tumors, including IFNα/γ responses, TNF/NF-κB and IL-6/JAK/STAT3 signaling (Fig. 3e). These inflammatory programs were largely attenuated at late stage and replaced by proliferative and metabolic signatures (Fig. 3e), indicating a shift toward tumor-supportive transcriptional programs during melanoma progression. Despite robust IFN responses in bulk early palpable tumors and sgTFAM YUMM1.7 cells, scRNA-seq revealed minimal innate immune activation within malignant cells at either stage, whereas immune cells, particularly myeloid populations, accounted for the predominant inflammatory and cytosolic DNA-sensing activity at the early timepoint (Fig. 3e-g). This result indicates a paracrine cGAS-STING and type I IFN signaling within the TME, whereby the early tumor-derived signals may induce epigenetic remodeling that sustain inflammatory programs in immune cells rather than tumor cell intrinsic persistent innate immune activation. Malignant cells in early TFAM^KD-MEL^ tumors exhibited metabolic stress, characterized by enrichment of mTORC1 signaling and hypoxia, indicating metabolic reprogramming toward glycolysis. Reclustering of the malignant compartment further resolved five transcriptional states corresponding to established melanoma phenotypes: melanocytic (*Pmel*, *Dct*), AP-1 stress-adaptive (*Nr4a2*, *Atf3*, *Zfp36*, *Egr2*), neural crest stem-like (*Ngfr*, *Axl*, *Lef1*, *Nes*), mesenchymal-transitory (*Fbn1*, *Wnt4*, *Elovl5*) and mesenchymal-invasive (*Zeb1*, *Postn*, *Vcan*, *Mmp2*) states (Extended Data Fig. 3c-e). Notably, the mesenchymal-invasive MEL5 population showed a transition from reduced to increased OXPHOS activity and strong enrichment of invasive programs relative to TFAM^WT-MEL^ controls (Extended Data Fig. 3f–h), linking the late metabolic shift to acquisition of an invasive malignant state.

**Fig. 3:**
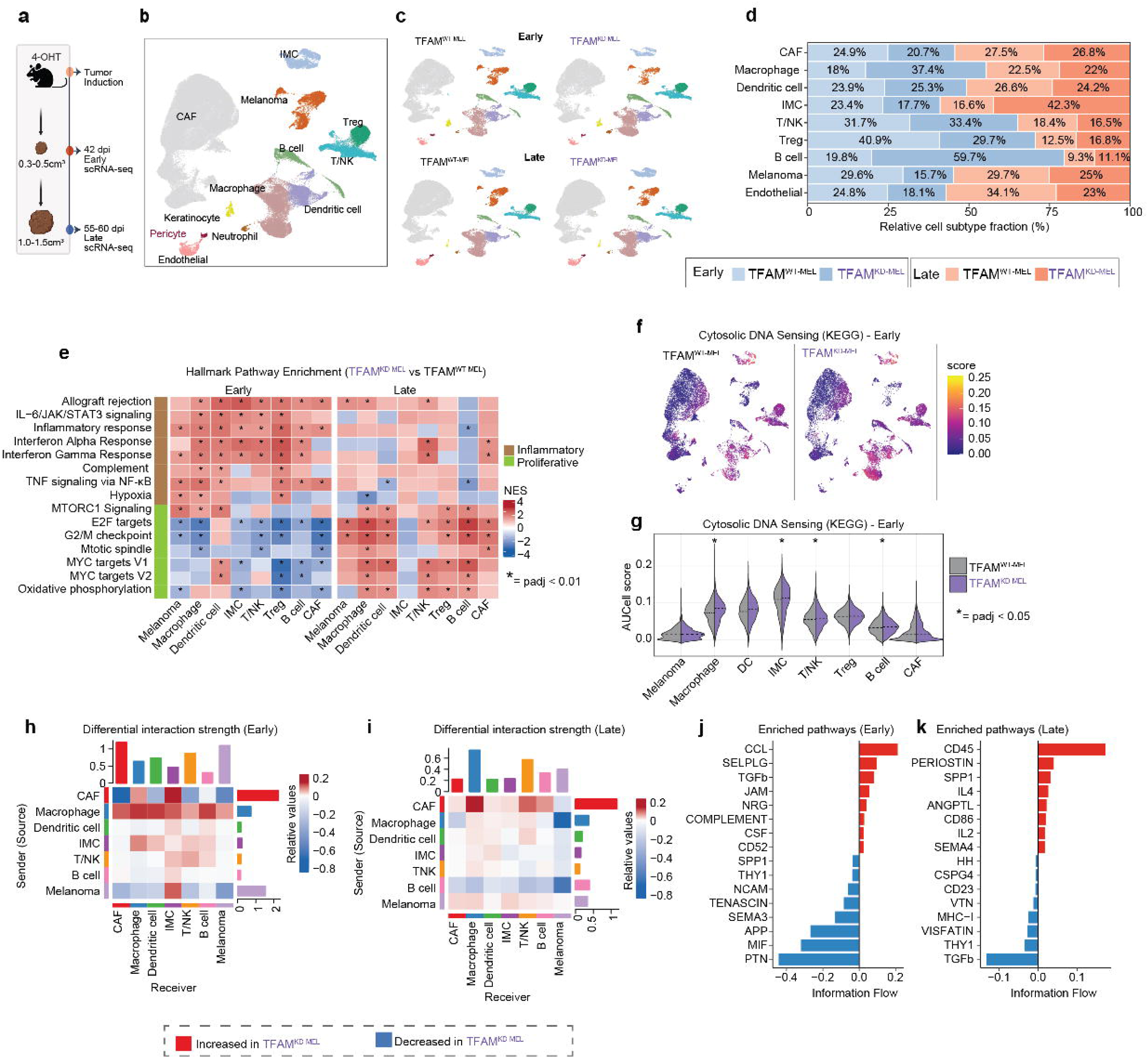
scRNA-seq reveals dynamic remodeling of the TME in TFAM^KD-MEL^ tumors. a, Schematic of the time-course scRNA-seq analysis. **b,c**, UMAP plots of integrated scRNA-seq data from early and late TFAM^KD-MEL^ and TFAM-^WT-MEL^ tumors, colored by major cell type (b) or split by genotype and time point (c). d, Stacked bar plot showing relative abundance of major cell types across TFAM^KD-MEL^ and TFAM^WT-MEL^ tumors. e, Hallmark pathway enrichment across indicated cell types at early and late stages. Single-cell counts were aggregated per sample and analyzed as pseudobulk using edgeR, followed by gene set enrichment analysis with fgsea. NES indicates enrichment of pathways in TFAM^KD-MEL^ (red) or TFAM^WT-MEL^ (blue) tumors. Selected pathways significantly enriched in at least two cell types are shown. Asterisks denote padj < 0.05. f, UMAP projections of KEGG Cytosolic DNA Sensing Pathway scores in TFAM^WT-MEL^ (left) and TFAM^KD-MEL^ (right) tumors at the early stage. g, Violin plots of the Cytosolic DNA Sensing Pathway signature scores for indicated cell types. Dashed lines indicate median scores. Statistical significance was assessed using the pseudobulk edgeR-fgsea approach described in e; Asterisks denote *padj* < 0.05. h,i, Heatmaps showing differential cell-cell communication strength inferred by CellChat at early (h) and late (i) stages. j,k, Signaling pathways with differential information flow between TFAM^KD-MEL^ and TFAM^WT-MEL^ tumors at early (j) and late (k) stages. Red and blue indicate increased and decreased information flow, respectively, in TFAM^KD-MEL^ tumors.

To map intercellular communication networks underlying TME remodeling, we performed CellChat analysis. TFAM^KD-MEL^ tumors displayed a marked increase in CAF-myeloid interactions with stage-dependent shifts in signaling hierarchy (Fig. 3h-k; Extended Data Fig. 4a, b). At early stage, macrophages represented the dominant signaling source, whereas at late-stage CAFs emerged as the principal signaling hub (Fig. 3h, i). Early TFAM^KD-MEL^ tumors were enriched for inflammatory and immune recruitment pathways including CCL, CSF, SELPLG, and complement signaling, whereas late-stage TFAM^KD-MEL^ tumors showed enrichment of immunoregulatory pathways including SPP1, IL4, and CD86 (Fig. 3j, k). Macrophage-to-CAF interactions via Il1b-Il1r1/Il1rap and Tgfb1-Tgfbr1/2 were enriched in early TFAM^KD-MEL^ tumors, suggesting that macrophage-derived inflammatory signals contribute to early CAF remodeling (Extended Data Fig. 4a, b). The CAF-derived Ccl2-Ccr2 axis was consistently enriched across both stages, supporting sustained monocyte recruitment to TFAM^KD-MEL^ tumors (Extended Data Fig. 4a, b). At late stage, CAF-derived Pros1-Axl, Gas6-Axl, and periostin-integrin (Itgav-Itgb5) interactions were significantly increased in macrophages, suggesting a role for CAFs in sustaining immunoregulatory macrophage states. Cxcl1-Cxcr2 signaling was also markedly elevated in TFAM^KD-MEL^ tumors, consistent with recruitment of Cxcr2-dependent myeloid populations. Melanoma cell-derived Lgals9 signaling through Cd44 and Havcr2, and Csf1-Csf1r signaling toward macrophages, were increased in early TFAM^KD-MEL^ tumors. The upregulation of Lgals9 and Csf1 in both early TFAM^KD-MEL^ tumors and sgTFAM YUMM1.7 cells implicates tumor cell-intrinsic mtDNA instability in driving myeloid recruitment and activation signals within the TME.

### mtDNA instability drives an invasion-permissive and immunoregulatory CAF ecosystem

Cancer-associated fibroblasts are key regulators of extracellular matrix organization and immune–stromal signaling within the TME. To define how tumor cell-intrinsic mtDNA instability influences the stromal compartment, we performed fibroblast subclustering analysis. Unbiased clustering combined with functional pathway scoring identified eight CAF subclusters (C0-C7) organized into six major subtypes (Fig. 4a, b; Extended Data Fig.5a-d), anchored to fibroblast states defined in recent single-cell and spatial atlases ^22,23^. Cluster C0 corresponded to a quiescent developmental CAF (dCAF) population characterized by *Hopx, Cd34, Reln,* and *Sox8*, whereas Cluster C1 represented an inflammatory CAF (iCAF) state defined by *Il6, Cxcl1, Cxcl2,* and *Ccl2* together with activation of NF-κB, TGF-β, and cytokine signaling pathways. Cluster C2 exhibited endothelial-associated features (*Cdh5, Cav1, Cav2*), consistent with an endothelial-to-mesenchymal transition CAF (EndMT-CAF) state, whereas Cluster C3 defined a myofibroblast CAF (myCAF) population expressing *Acta2, Tagln, Postn, Lrrc15,* and *Mmp11*. Cluster C4 displayed a vascular CAF (vCAF) program marked by *Rgs5, Pdgfrb,* and *Notch3* together with pro-angiogenic mediators including *Gdf10, Angpt2,* and *Pdgfd*. Cluster C5 represented a stress-associated EndMT-CAF state expressing *Rasd1, Rgs16, Slc2a1,* and *Icam1* and enriched for hypoxia, TNF/NF-κB, IL-6/JAK/STAT3, and IFN signaling programs. Cluster C6 corresponded to a *Pi16* fibroblast progenitor population (*Pi16, Dpp4, Dpt, Sfrp2,* and *Sfrp4*) enriched for IFN-response pathways, whereas Cluster C7 represented a proliferating (*Mki67*, *Top2a)* CAF state (Extended Data Fig.5d). Notably, dCAFs and iCAFs exhibited substantial transcriptional overlaps, consistent with a continuum between resting and inflammatory fibroblast states, and iCAFs were preferentially enriched in TFAM^KD-MEL^ tumors, consistent with inflammatory CAF programs associated with advanced melanoma ^23^.

**Fig. 4:**
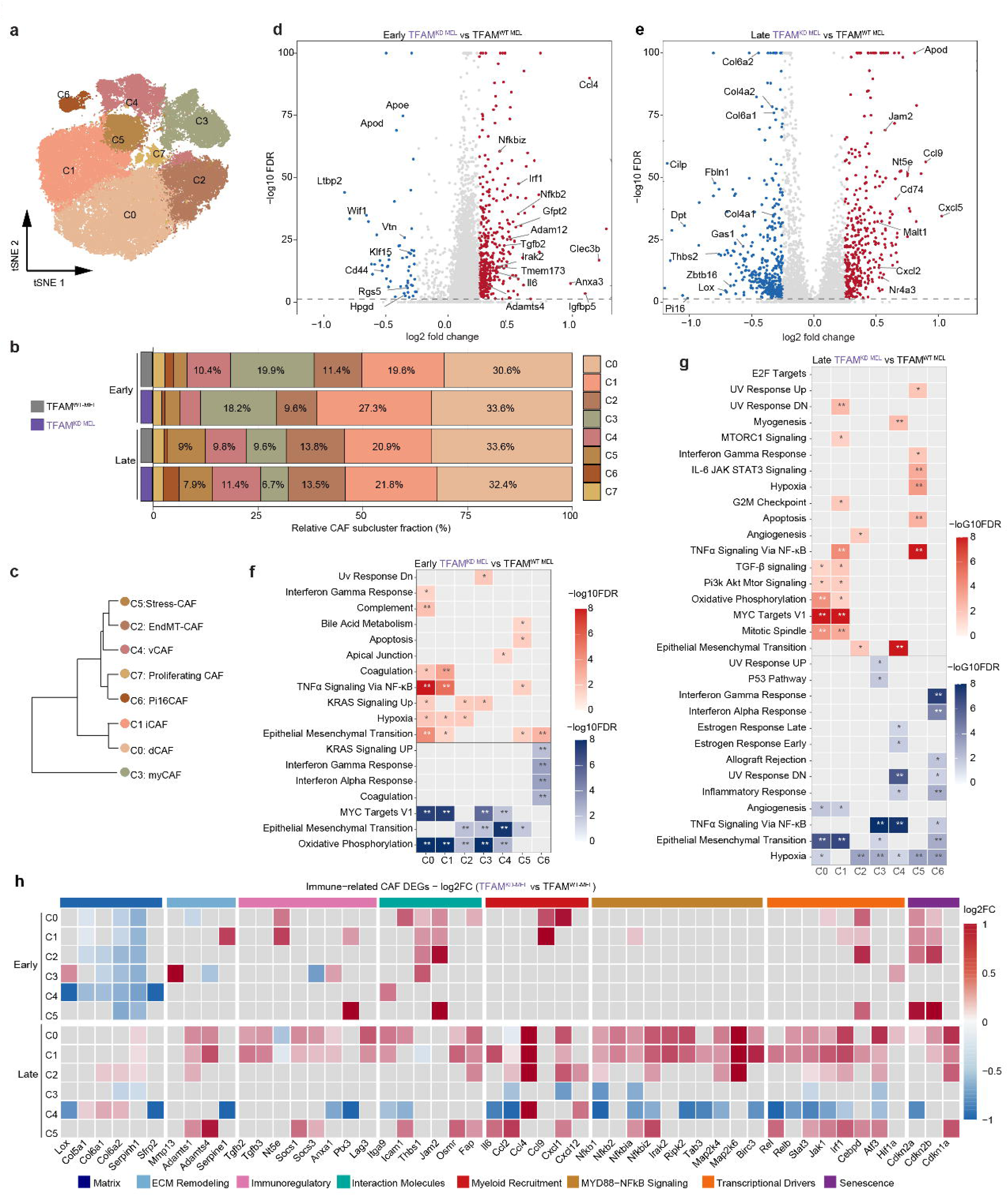
CAFs in TFAM^KD-MEL^ tumors acquire an immunomodulatory phenotype. a, UMAP plot of CAF subclusters from TFAM^KD-MEL^ and TFAM^WT-MEL^ tumors. b, Stacked bar plot showing relative abundance of CAF subclusters in TFAM^KD-MEL^ and TFAM^WT-MEL^ tumors at early and late time points. c, Hierarchical clustering dendrogram showing transcriptomic relationships among CAF subclusters. d, e, Volcano plots showing DEGs between TFAM^KD-MEL^ and TFAM^WT-MEL^ CAFs at early (d) and late (e) time points. CAF subclusters were aggregated by genotype within each time point; selected extracellular matrix, inflammatory, and immunoregulatory genes are highlighted. f, g, Heatmaps showing Hallmark pathway enrichment among DEGs in TFAM^KD-MEL^ relative to TFAM^WT-MEL^ CAFs at early (f) and late (g) time points. Red indicates pathways upregulated and blue indicates downregulated in TFAM^KD-MEL^ CAFs. Statistical significance was assessed using Benjamini–Hochberg correction (*FDR < 0.05, **FDR < 0.01). h, Heatmap of matrix- and immune-related DEGs (FDR < 0.05) in TFAM^KD-MEL^ versus TFAM^WT-MEL^ CAFs, grouped into functional modules. Color indicates log□ fold change; non-significant genes are shown in white.

To define mtDNA instability-associated transcriptional changes in CAFs, subclusters were aggregated by genotype and subjected to differential expression analysis. CAFs from TFAM^KD-MEL^ tumors showed reduced expression of matrix organization and stromal homeostatic genes (*Col6a1*, *Col6a2*, *Fbln1*, *Dpt*) and higher expression of inflammatory mediators (*Il6*, *Cxcl2*, *Ccl4*), immunomodulatory genes (*Nt5e*, *Cd74*), and NF-κB-associated regulators (*Nfkbiz*, *Nfkb2*, *Irf1*) (Fig. 4d, e), indicating acquisition of inflammatory and immunoregulatory programs with concurrent loss of structural matrix integrity relative to TFAM^WT-MEL^ tumors.

Cluster-level analysis revealed CAF state-specific remodeling in TFAM^KD-MEL^ tumors (Fig. 4f-h). At early stage, the predominant stromal response was broad suppression of OXPHOS and MYC target programs across structural CAF subtypes, and reduced expression of genes involved in collagen assembly and matrix organization (*Lox*, *Col6a1*, *Col4a1*, *Col4a2*), consistent with loss of matrix integrity relative to TFAM^WT-MEL^ tumors and resembling invasion-permissive CAF states described in age-associated melanoma (Kaur et al., 2019). Inflammatory signaling was enriched in dCAF (C0) and iCAF (C1) in TFAM^KD-MEL^ tumors, including TNF/NF-κB and hypoxia-associated programs. At late stage, CAFs in TFAM^KD-MEL^ tumors showed further enrichment of immunoregulatory transcriptional programs (Fig. 4f-h). dCAF (C0), iCAF (C1), and stress-CAF (C5) exhibited higher enrichment of immunoregulatory signaling including TNFα/NF-κB, TGF-β, and IL-6/JAK/STAT3 pathways, and higher expression of genes involved in immunoregulation (*Tgfb2/3*, *Nt5e*, *Socs1-3*), myeloid recruitment (*Ccl2*, *Ccl9*, *Cxcl1*), NF-κB pathway components (*Nfkbiz*, *Irak2*), and inflammatory transcription factors (*Irf1*, *Stat3*), overlapping with inflammatory CAF state associated with melanoma progression and immunotherapy resistance ^23^.

Pi16□ fibroblast progenitors (C6) exhibited downregulation of IFN-response, antigen-presentation, and inflammatory pathways in TFAM^KD-MEL^ tumors relative to TFAM^WT-MEL^ tumors at both stages (Fig. 4e–h; Extended Data Fig. 5e). Given the association of Pi16□ fibroblasts with immune-interactive stromal niches and favorable prognosis ^22^, these findings indicate that mtDNA instability impairs immune-supportive fibroblast progenitor compartments within the tumor stroma. In summary, mtDNA instability drives stromal remodeling characterized by reduced expression of genes involved in matrix integrity, alongside progressive amplification of chronic TNFα/NF-κB and IL-6/JAK/STAT3 engagement and immunoregulatory transcriptional programs.

### mtDNA instability drives macrophage infiltration and tolerogenic reprogramming

To define mtDNA instability–associated changes in tumor-infiltrating immune cells, we subclustered CD45^+^ immune cells and identified diverse lymphoid and myeloid populations (Extended Data Fig. 6a, b). Lymphoid cells comprised T cells (*Cd3e*, *Trac*), regulatory T cells (*Ctla4*, *Foxp3*), natural killer cells (NK: *Ncr1*, *Klrb1c*), innate lymphoid cells (ILC: *Il17rb, Il5*), gamma-delta T cells (γδT: *Trdc*, *Trdv4*), and B cell populations (*Cd19*) spanning progenitor (*Rag1*, *Igll1*) and mature (*Cxcr5*, *Ms4a1*) states. Myeloid cells represented the dominant immune compartment and included three Adgre1^+^Csf1r^+^ macrophage populations, conventional dendritic cells (cDC1: *Xcr1*, *Clec9a*; cDC2: *Cd209a*, *Clec10a*), plasmacytoid DCs (*Siglech*, *Ccr9*), Langerhans cells (LC: *Epcam*, *Cd207*), and mature migratory dendritic cells (*Mreg*, *Fscn1*) (Extended Data Fig. 6d). Early TFAM^KD-MEL^ tumors exhibited increased macrophage infiltration, particularly of *Chil3*^+^*Hp*^+^ inflammatory macrophages (Mac_1), and reduced Treg abundance relative to TFAM^WT-MEL^ controls (Extended Data Fig. 6c). At late stage, Treg frequency within the CD3^+^ compartment was increased in TFAM^KD-MEL^ tumors (Extended Data Fig. 6l). Multicolor flow cytometry independently demonstrated increased Treg abundance in late TFAM^KD-MEL^ tumors relative to TFAM^WT-MEL^ controls (Extended Data Fig. 7d).

Within the Adgre1^+^Ly6c2+ myeloid compartment negative for Csf1r, we identified a transcriptionally distinct myeloid population that expanded markedly in late TFAM^KD-MEL^ tumors (Extended Data Fig. 6a, b). This population expressed genes associated with both monocytic (*Adgre1*, *Itgam*, *Ly6c2*) and granulocytic (*Fcer1a*, *Gata2*, *Hdc*) myeloid lineage markers, which we annotated as immature myeloid cells (IMC). Reactome pathway activity scoring revealed that IMCs exhibited the highest activity for cytosolic DNA sensing and autophagy, type 2 cytokine and IL-6 family signaling, arachidonic acid metabolism, Fc receptor-mediated calcium mobilization, and matrix metalloproteinase activation (Extended Data Fig. 6e), indicating a functionally distinct myeloid state with broad innate immune signaling and tissue-remodeling capacity. IMCs further showed enrichment of lipid mediator synthesis genes (*Alox5*, *Ptgs2*), histamine biosynthesis (*Hdc*), immune checkpoint ligands (*Cd274*, *Pdcd1lg2*), inhibitory receptors (*Lilrb4a*, *Lilrb4b*), and the matrix remodeling gene *Mmp9*. Relative to TFAM^WT-MEL^ tumors, IMCs from TFAM^KD-MEL^ tumors showed higher expression of inflammatory and immunoregulatory mediators (Extended Data Fig. 6i). To further characterize this population, we performed MDSC signature scoring using previously defined gene sets ^24^. IMCs exhibited significantly higher MDSC signature scores than other myeloid populations, with further enrichment in TFAM^KD-MEL^ tumors (Extended Data Fig. 6f, g). Collectively, mtDNA instability-associated inflammatory TME promotes enrichment of an MDSC-like myeloid population with inflammatory and immunoregulatory features.

Macrophages represented the most abundant immune population in TFAM^KD-MEL^ tumors and are established regulators of tumor progression and immune evasion in melanoma ^25–27^. To investigate how mtDNA instability shapes macrophage states within the TME, we performed high-resolution subclustering of the Adgre1^+^Csf1r^+^ macrophage compartment, and identified six subclusters (Fig. 5a, b). MoMac cells expressed *Ly6c2*, *Vcan*, *Plac8* and *Chil3*, consistent with a monocyte-derived transcriptional state. ISG^+^ Mac shared MoMac marker genes but was distinguished by strong enrichment of ISGs including *Isg15* and *Mx1*. Marchf1+ Mac co-expressed *Marchf1*, which promotes MHC-II ubiquitination and lysosomal degradation ^28^, and *Zeb2*, a regulator of tumor associated macrophage (TAM) differentiation and function ^29^, while lacking clear subtype-specific identity markers, consistent with an intermediate macrophage state. *Trem2*+ Mac displayed a mature macrophage program marked by *Trem2*, *Apoe*, *Mrc1*, and *C1qb*, together with efferocytosis-associated genes *Gas6*, *Stab1*, and *Mertk*. *Retnla*+ Mac selectively expressed *Retnla*, *Cd226*, *Mgl2*, *Fcrls*, and lipid-handling genes *Lpl* and *Pltp*, consistent with an alternatively activated macrophage state. Ace+ Mac, a small subpopulation, was distinguished by non-classical monocyte and lipid metabolism-associated genes including *Ace*, *Treml4*, and *Fabp4*. Reactome pathway analysis confirmed distinct functional programs across macrophage states (Extended Data Fig. 6e). Macrophage subset composition analysis revealed increased MoMac frequency (54.2% versus 43.0%) and decreased *Retnla*+ Mac frequency (6.5% versus 17.0%) in early TFAM^KD-MEL^ tumors relative to TFAM^WT-MEL^ tumors (Fig. 5c). At late stage, the MoMac compartment contracted (35.6% versus 44.2%) while Trem2+ Mac expanded (45.1% versus 38.3%), indicating a shift toward immunoregulatory TAM states. Monocle3 pseudotime analysis, initiated from *Ly6c2*/*Sell*/*Vcan*-high macrophages, identified a transcriptional continuum from inflammatory monocyte-derived states to mature TAM states, along which inflammatory mediators (e.g., *Il1b*, *Hif1a*) declined and immunoregulatory TAM markers (e.g., *Mrc1*, *Spp1*) increased (Fig. 5d, e). TFAM^KD-MEL^ macrophages showed more pronounced *Mrc1* and *Spp1* expression differences along this trajectory (Fig. 5e). Flow cytometric analysis of late-stage tumors demonstrated an increased frequency of CD206^+^ macrophages, higher CD206 expression levels, and reduced Ly6C^+^ macrophage abundance in TFAM^KD-MEL^ tumors (Fig. 5f, g), independently supporting enrichment of an immunoregulatory TAM at late stage.

**Fig. 5:**
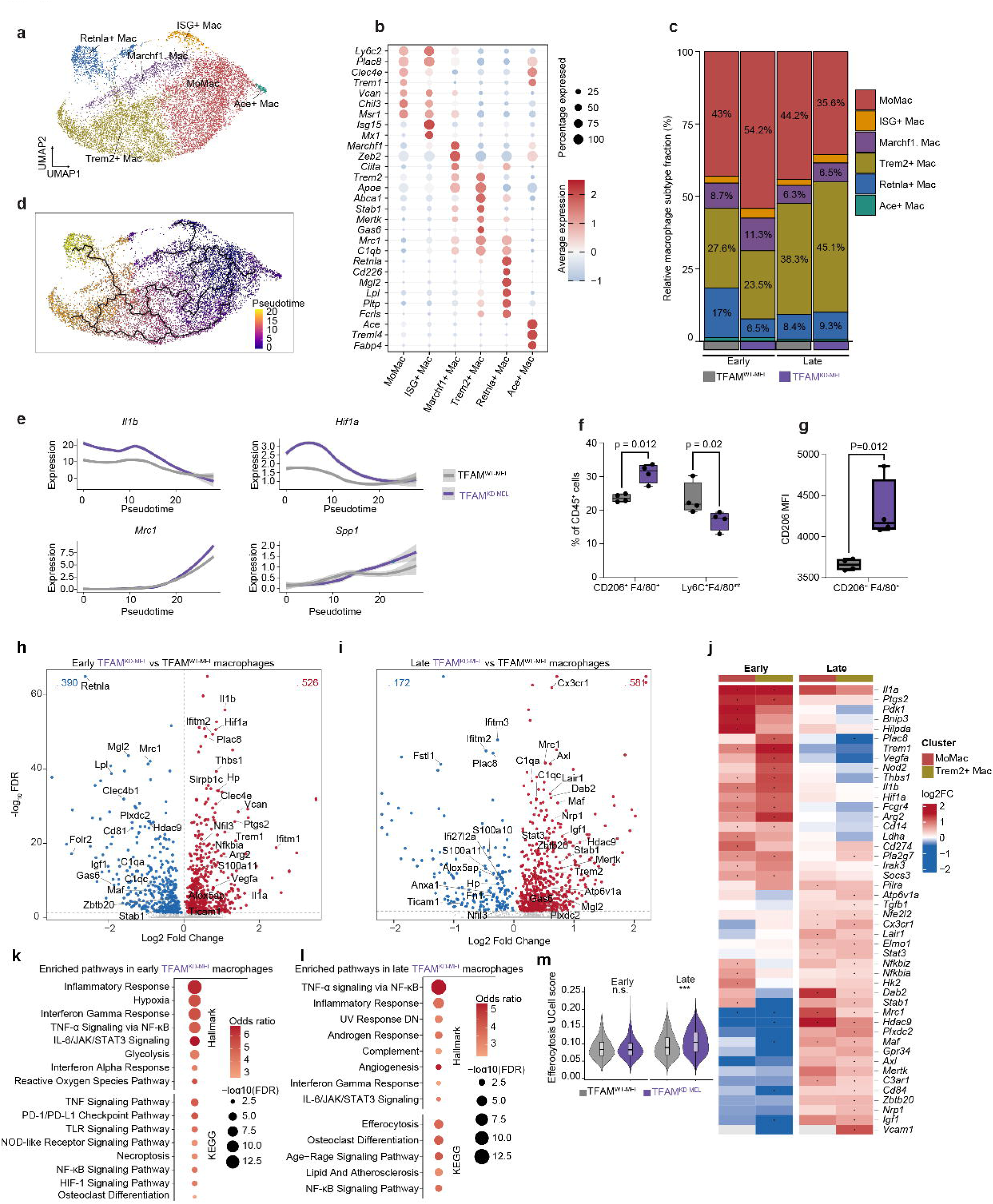
Tumor cell-intrinsic mtDNA instability drives temporal macrophage reprogramming. a, UMAP plot of macrophage subclusters in TFAM^KD-MEL^ and TFAM^WT-MEL^ tumors, colored by cluster. b, Dot plot showing marker gene expression across macrophage subclusters. Dot size indicates the percentage of expressing cells, and color represents scaled average expression. c, Stacked bar plot showing the relative abundance of macrophage subclusters in TFAM^KD-MEL^ and TFAM^WT-MEL^ tumors at early and late stages. d, Pseudotime trajectory of macrophage subclusters inferred using Monocle3. e, Smoothed gene expression dynamics along the pseudotime trajectory for representative inflammatory (*Il1b*, *Hif1a*) and immunoregulatory (*Mrc1*, *Spp1*) markers in TFAM^KD-MEL^ and TFAM^WT-MEL^ tumors. f,g, Flow cytometric analysis of late-stage tumors showing increased CD206□ macrophage frequency (f) and CD206 expression (g), together with reduced Ly6C^hi^F4/80^int^ inflammatory macrophages in TFAM^KD-MEL^ tumors. h,i, Volcano plots showing DEGs between TFAM^KD-MEL^ and TFAM^WT-MEL^ macrophages at early (h) and late (i) stages. Genes enriched in TFAM^WT-MEL^ TAMs (left) and TFAM^KD-MEL^ TAMs (right) are shown, with selected genes with established pro- or antitumor functions highlighted. Significant genes (FDR < 0.05, |log□ FC| > 0.25) are highlighted in red (upregulated in TFAM^KD-MEL^) or blue (downregulated in TFAM^KD-MEL^ relative to TFAM^WT-MEL^). j, Heatmap showing log□ fold change of selected inflammatory mediators (upregulated early) and tolerogenic genes (upregulated late) (FDR < 0.05, |log□ FC| > 0.25) in Trem2□ and MoMac TFAM^KD-MEL^ macrophages relative to TFAM^WT-MEL^ macrophages. k,l, Hallmark and KEGG pathway enrichment of significantly upregulated DEGs in TFAM^KD-MEL^ macrophages (FDR < 0.05). m, Violin plots showing single-cell-level scores for an efferocytosis-related gene signature in macrophages at early and late stages. Asterisks denote adjusted *P* < 0.05; ns, not significant.

To define the molecular features underlying macrophage remodeling, we performed differential expression analysis of macrophages aggregated by genotype and tumor stage. In early TFAM^KD-MEL^ tumors, macrophages showed higher expression of inflammatory mediators (e.g., *Il1b*, *Trem1*, *Ptgs2*), hypoxia-associated and angiogenic factors (e.g., *Vegfa*, *Hif1a*), and matrix-remodeling genes (e.g., *Thbs1*, *Vcan*, *Tgfbi*), together with reduced expression of tissue-adaptive macrophage genes (*C1qa, Mrc1*, *Igf1*, *Axl*, *Gas6*, *Stab1*) relative to TFAM^WT-MEL^ tumors (Fig. 5h), indicating an inflammatory macrophage state with angiogenic and matrix-remodeling features. Pathway enrichment corroborated these transcriptional changes, with early TFAM^KD-MEL^ macrophages enriched for IFN responses, IL-6/JAK/STAT3, TNF/NF-κB, hypoxia, and glycolysis pathways, consistent with a metabolically rewired inflammatory state (Fig. 5k). At late stage, macrophages in TFAM^KD-MEL^ tumors showed increased tissue-adaptive transcriptional program relative to TFAM^WT-MEL^ tumors, characterized by increased expression of immunoregulatory transcription factors (*Maf*, *Smad3*, *Nfe2l2*, *Zbtb20*), inhibitory receptors (*Trem2*, *Nrp1*, *Lair1*, *Cd84*), and genes associated with scavenging and efferocytic function (*Mertk*, *Gas6*, *Stab1*, *Dab2*, *Axl*, *Mrc1*), together with genes involved in phagosomal and lysosomal function (*Elmo1*, *Pik3c3*, *Atp6v1a*) (Fig. 5i). Pathway analysis further identified enrichment of efferocytosis, endocytosis, lipid metabolism, and adhesion pathways, consistent with a tissue-remodeling macrophage program (Fig. 5i). Efferocytosis-related gene signature scoring independently demonstrated enrichment in macrophages from late TFAM^KD-MEL^ tumors (Fig. 5m), consistent with mtDNA instability-driven acquisition of a tissue-remodeling efferocytic macrophage state linked to tumor progression and resistance to ICB ^25,30^.

Cluster-level differential expression analysis revealed that at an early stage, mtDNA instability broadly shifted macrophages toward inflammatory activation, with MoMac cells exhibiting the most extensive inflammatory reprogramming (Fig. 5j; Table S3). MoMac cells showed higher expression of glycolytic and hypoxia-adaptation genes (*Hif1a*, *Bnip3*, *Hilpda*, *Ldha*, *Pdk1*, *Hk2*), NF-κB pathway components (*Nfkbia*, *Nfkbiz*), and inflammatory mediators (*Il1a*, *Trem1*, *Ptgs2*) relative to TFAM^WT-MEL^ tumors, defining a metabolically rewired inflammatory state (Fig. 5j). Trem2+ Mac showed higher expression of inflammatory mediators (*Il1b*, *Nos2*), angiogenic and matrix-remodeling genes (*Vegfa*, *Thbs1*), and immunoregulatory effectors (*Socs3*, *Arg2*, *Fcgr4*) in early TFAM^KD-MEL^ tumors. *Retnla*+ macrophages, markedly reduced in early TFAM^KD-MEL^ tumors, showed loss of canonical identity markers (*Retnla*, *Ear2*, *Fcrls*) and induction of inflammatory and tissue-remodeling genes (*Il1b*, *Trem1*, *Ptgs2*, *Vegfa*, *Thbs1*), indicating mtDNA instability-driven displacement of tissue-adaptive macrophage identity. At late stage, Trem2+ Mac displayed the strongest enrichment of immunoregulatory (*Cx3cr1*, *Stat3*, *Nrp1*, *Hdac9*, *Smad3*, *Cd84*, *Prkaa1*, *C3ar1*, *Pilra*) and efferocytosis-related genes (*Mertk*, *Stab1*, *Axl*, *Elmo1*, *Dab2*, *Atp6v1a*) in TFAM^KD-MEL^ tumors relative to TFAM^WT-MEL^ tumors, identifying this subset as the principal macrophage population driving the immunoregulatory program at late stage.

These findings collectively suggest that tumor cell-intrinsic mtDNA instability drives inflammatory macrophage expansion and temporal reprogramming toward an efferocytosis-associated immunoregulatory state, positioning macrophages as a key population through which mtDNA instability shapes the immune landscape of melanoma.

### TFAM^KD-MEL^ macrophage signatures stratify survival and therapeutic response in human melanoma

To determine whether TFAM^KD-MEL^ macrophage signatures map to clinically relevant myeloid states in human melanoma, we subsetted monocyte and macrophage populations from a meta-analysis of melanoma scRNA-seq datasets from patients treated with ICB ^31^ and performed unsupervised re-clustering and annotation. Six subpopulations were identified, including FCN1+ monocytes, neutrophils, and four macrophage states (IFN+, SPP1+, MRC1/C1Q+, and APOE/C1Q+ Mac; Fig. 6a, b). FCN1+ Mono was characterized by monocyte-associated genes including *FCN1*, *PLAC8*, and *VCAN*, whereas IFN+ Mac expressed IFN-inducible genes including *CXCL9*, *ISG15*, *IDO1*, *LGALS2*, and *SLAMF7*. SPP1+ Mac was defined by expression of *SPP1*, *INHBA*, *GPNMB*, and *FABP5*. A population originally annotated as monocytes based on *S100A8* and *S100A9* expression was re-annotated as neutrophils based on expression of the neutrophil-specific markers *FCGR3B* and *CSF3R*. The population originally classified as M2 macrophages was resolved into two transcriptionally distinct states: APOE+ Mac, characterized by higher *APOE* and *APOC1* expression, and MRC1+ Mac, distinguished by expression of *MRC1*, *STAB1*, and *MERTK*, markers associated with scavenging and efferocytic function.

**Fig. 6:**
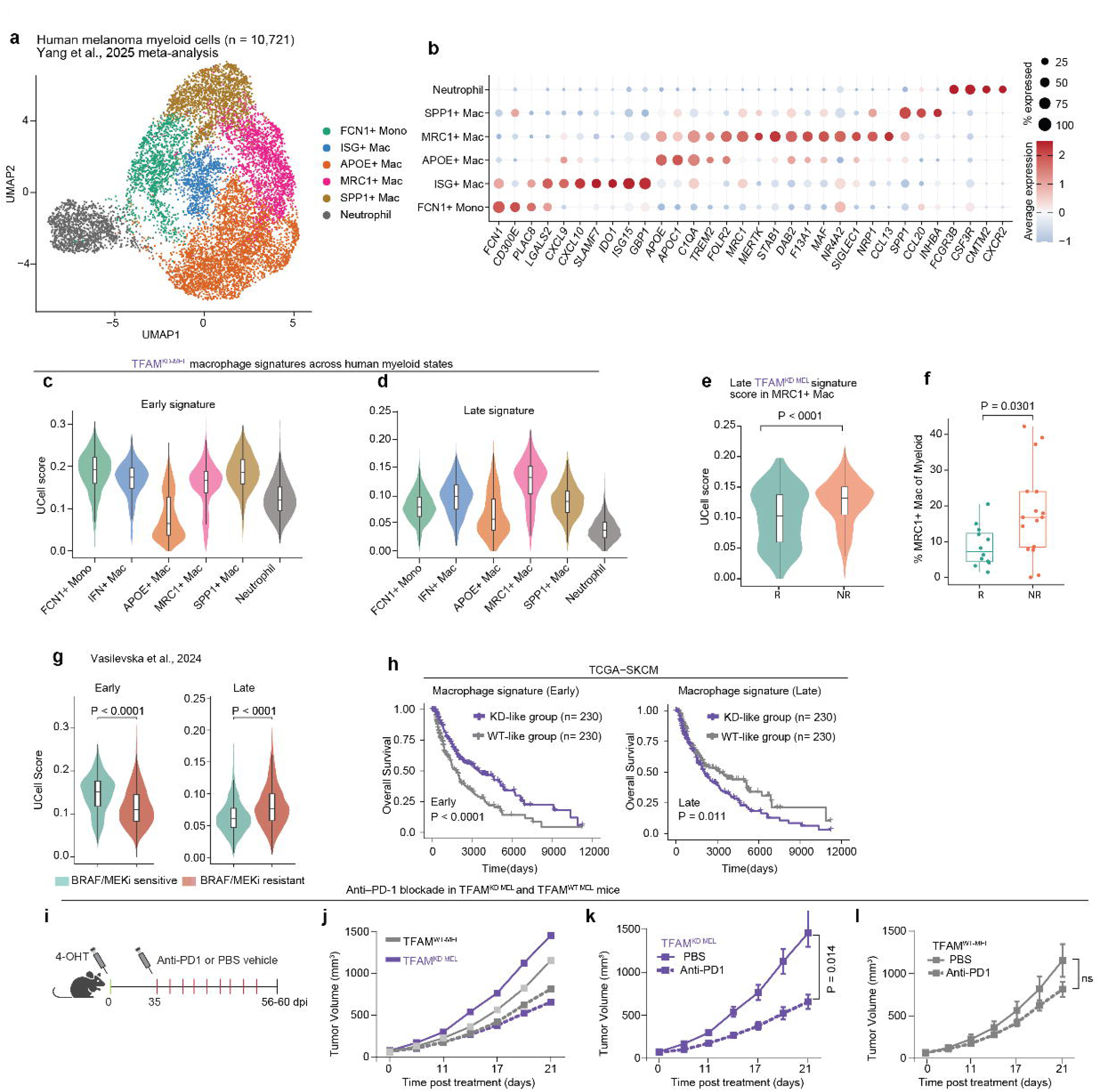
TFAM^KD-MEL^ macrophage signatures are conserved in human melanoma and predict clinical outcomes and therapy response. **a**, UMAP plot of macrophages, monocytes and neutrophils from a meta-analysis of melanoma scRNA-seq datasets of patients treated with ICB ^31^. b, Dot plot shows marker gene expression defining the cluster identities in a. Dot size indicates the percentage of cells expressing each gene, and color indicates scaled average expression. c,d, Violin plots show enrichment of the early (c) and late (d) TFAM^KD-MEL^ macrophage signatures in each cluster. e, Late TFAM^KD-MEL^ macrophage signature scores in MRC1□ macrophages from ICB responders and non-responders (*P* < 0.0001, two-sided Wilcoxon rank-sum test). f, Frequency of MRC1□ macrophages among myeloid cells in ICB responders and non-responders (n = 29; *P* = 0.030, two-sided Wilcoxon rank-sum test). g, Enrichment of the early (left) and late (right) TFAM^KD-MEL^ macrophage signatures in macrophages from BRAF/MEK inhibitor-sensitive and -resistant tumors (Vasilevska et al., 2024). h, Kaplan–Meier overall survival curves for the TCGA-SKCM cohort, stratified at the median into high (KD-like) and low (WT-like) signature-scoring groups based on the early (left) and late (right) TFAM^KD-MEL^ macrophage signatures. i, Experimental design for anti-PD-1 treatment of TFAM^KD-MEL^ and TFAM^WT-MEL^ tumor-bearing mice. j, Tumor growth curves comparing TFAM^WT-MEL^ and TFAM^KD-MEL^ tumors (n = 6–9 mice per group). k, Tumor growth curves of TFAM^KD-MEL^ tumors treated with anti-PD-1 antibody or PBS vehicle control. l, Tumor growth curves of TFAM^WT-MEL^ tumors treated with anti-PD-1 antibody or PBS vehicle control. Statistical significance was assessed using a two-sided Wilcoxon rank-sum test with Benjamini– Hochberg correction (e,f), a two-sided log-rank test (h), and mixed-effects analysis (j–l). Adjusted *P* values are reported for e,f.

We next mapped the top-ranked genes upregulated in TFAM^KD-MEL^ macrophages relative to TFAM^WT-MEL^ controls onto these human myeloid subtypes. The early signature was enriched in FCN1□ Mono and SPP1□ Mac, consistent with an inflammatory and angiogenic transcriptional program (Fig. 6c). In contrast, the late TFAM^KD-MEL^ macrophage signature was selectively enriched in MRC1□ Mac (Fig. 6d), which expressed genes associated with tissue-adaptive macrophage states, efferocytosis and immune regulation, including *MRC1*, *STAB1*, *MERTK*, *DAB2*, *F13A1*, *MAF*, *NR4A2*, *SIGLEC1*, *NRP1*, *PLTP*, and *CCL13*. Notably, the frequency of MRC1□ Mac, but not other subpopulations, was significantly higher in ICB non-responders than responders (n = 29; 12 responders, 17 non-responders; P = 0.030, Fig. 6f), and the late TFAM^KD-MEL^ macrophage signature was significantly enriched within MRC1 Mac from non-responders relative to responders (P < 0.0001, Fig. 6e). These findings link the late TFAM^KD-MEL^ macrophage program to an immunoregulatory MRC1 Mac state associated with ICB non-response in human melanoma.

The temporal macrophage programs were also associated with response to targeted therapy. In an independent cohort of melanoma patients treated with BRAF/MEK inhibitors ^27^, the early TFAM^KD-MEL^ macrophage signature was strongly enriched in macrophages from therapy-sensitive tumors (P < 0.0001), whereas the late signature was preferentially enriched in macrophages from therapy-resistant tumors (P < 0.0001, Fig. 6g). Together with the ICB response data, these findings show that the stage-dependent macrophage programs identified in TFAM^KD-MEL^ tumors, an early inflammatory and a late efferocytosis-associated, immunoregulatory programs, correspond to macrophage states associated with treatment response and resistance in human melanoma.

We next examined whether these temporal macrophage programs were associated with clinical outcome in human melanoma. In the TCGA-SKCM cohort, early and late TFAM^KD-MEL^ macrophage signature scores were calculated based on the relative enrichment of genes upregulated versus downregulated in TFAM^KD-MEL^ macrophages. Higher early TFAM^KD-MEL^ macrophage signature scores were associated with favorable overall survival (P < 0.0001, log-rank test), whereas higher late TFAM^KD-MEL^ macrophage signature scores were associated with poor overall survival (P = 0.011, log-rank test) (Fig. 6h). Thus, the temporal macrophage programs identified in TFAM^KD-MEL^ tumors capture distinct clinical associations in human melanoma, with the early program associated with longer survival and the late program with shorter survival.

To test whether the early mtDNA instability-associated IFN response confers sensitivity to immune checkpoint blockade, we treated mice bearing TFAM^KD-MEL^ or TFAM^WT-MEL^ tumors (50-100 mm³) with anti-PD-1 antibody or PBS vehicle control (Fig. 6i-l). Anti-PD-1 treatment significantly delayed tumor growth in TFAM^KD-MEL^ mice relative to vehicle-treated controls, whereas no significant effect on tumor growth was observed in TFAM^WT-MEL^ mice. Anti-PD-1 treatment did not abrogate tumor growth or induce sustained tumor control in TFAM^KD-MEL^ tumors, indicating a partial and transient response. Given the relatively T cell-poor nature of this model, the limited therapeutic effect may reflect insufficient T cell availability or resistance mechanisms within the TME.

Collectively, these findings establish that the macrophage transcriptional programs driven by tumor cell-intrinsic mtDNA instability are clinically stratified, with early inflammatory signatures mapping to human melanoma macrophage states associated with favorable survival and ICB response, whereas late efferocytic and immunoregulatory signatures map to states associated with poor prognosis and therapy resistance.

### cGAS is required for mtDNA stress-driven TME remodeling and melanoma progression

Our findings establish that melanoma cell-intrinsic mtDNA instability drives an inflammatory TME that promotes tumor progression and metastasis. Given that aberrant mtDNA release is a potent cGAS agonist ^3^ and that chronic cGAS-STING activation engages immunosuppressive programs associated with poor clinical outcomes ^13,14^, we tested whether cGAS is causally required for mtDNA instability-driven melanoma phenotypes by generating TFAM^KD-MEL^ mice in a cGAS-deficient background (TFAM^KD-MEL^cGAS^KO^). cGAS ablation reversed both the accelerated tumor growth and enhanced metastatic spread observed in TFAM^KD-MEL^cGAS^WT^ mice (Fig. 7a, b). cGAS deletion had no effect on tumor growth in the absence of Tfam knockdown, as TFAM^WT-MEL^cGAS^KO^ and TFAM^WT-MEL^cGAS^WT^ mice displayed similar tumor growth kinetics (Fig. 7a), establishing that cGAS is required for mtDNA instability-driven tumor promotion but dispensable for baseline melanoma growth. Immunohistochemical analysis demonstrated that cGAS deletion abrogated the enhanced lymph node metastasis observed in TFAM^KD-MEL^cGAS^WT^ tumors. TFAM^KD-MEL^cGAS^KO^ mice exhibited significantly fewer SOX10^+^ melanoma cells in tumor-draining inguinal lymph nodes (Fig. 7f) and reduced secondary cutaneous tumor burden relative to TFAM^KD-MEL^cGAS^WT^ mice (Fig. 7b), while metastatic incidence was unaffected by cGAS status in TFAM^WT-MEL^ mice, confirming that cGAS engagement is specific to the mtDNA instability context.

**Fig. 7:**
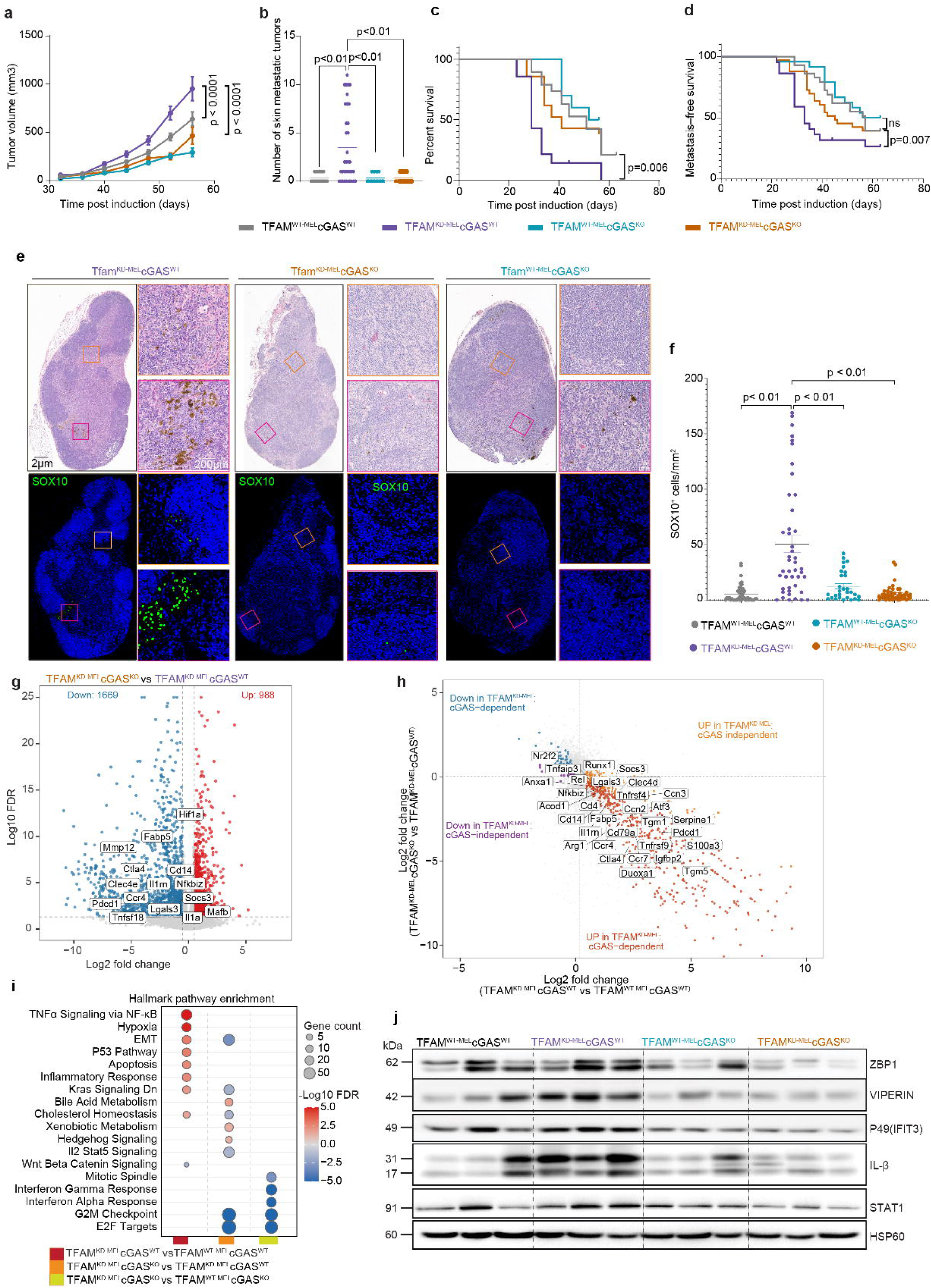
cGAS deficiency attenuates tumor growth and inflammatory signatures of TFAM^KD-MEL^ tumors. a, Tumor growth curves for TFAM^WT-MEL^cGAS^WT^, TFAM^KD-MEL^cGAS^WT^, TFAM^WT-MEL^cGAS^KO,^ and TFAM^KD-MEL^cGAS^KO^ mice (n = 50-60 per genotype); statistical significance assessed by two-way ANOVA with multiple comparisons correction. b, Quantification of macroscopic skin metastatic tumor nodules across groups. c, d, Kaplan-Meier plots showing overall survival from palpable tumor detection to humane endpoint (c) and metastasis-free survival (d). Statistical differences between all groups were assessed by log-rank (Mantel-Cox) test. e, Representative H&E-stained sections (upper panels) and immunofluorescence for SOX10 (green) with DAPI nuclear counterstain (blue; lower panels) of tumor-draining inguinal lymph nodes (ILNs) from all four genotypes (n = 25-30 per genotype). Scale bar, 2 μm and 200 μm. f, Quantification of SOX10^+^ cells in ILNs across groups; statistical significance assessed by two-way ANOVA with multiple comparisons correction. g, Volcano plots of DEGs in TFAM^KD-MEL^cGAS^KO^ versus TFAM^KD-MEL^cGAS^WT^. Significantly upregulated and downregulated genes (FDR < 0.05, |log2FC| > 0.5) are highlighted in red and blue, respectively; selected immune related genes are labeled. h, Scatter plot showing the transcriptional relationship between TFAM ablation and cGAS dependency in melanoma. The x-axis shows differential expressions between TFAM^KD-MEL^cGAS^WT^ and TFAM^WT-MEL^cGAS^WT^ tumors, and the y-axis shows the effect of cGAS ablation in TFAM^KD-MEL^ tumors (TFAM^KD-MEL^cGAS^KO^ versus TFAM^KD-MEL^cGAS^WT^). Genes are classified as cGAS-dependent if significantly altered in both comparisons with opposite directionality (FDR < 0.05, |log2FC| > 0.5, DESeq2), including genes induced by TFAM ablation and rescued by cGAS deletion (red; lower-right quadrant) and genes suppressed by TFAM ablation and restored by cGAS deletion (blue; upper-left quadrant). Genes regulated by TFAM independently of cGAS are shown in orange and purple. Selected cGAS-dependent genes are labeled. i, Volcano plots showing DEGs between TFAM^KD-MEL^cGAS^KO^ versus TFAM^KD-MEL^cGAS^W^. j, Significantly enriched Hallmark pathways (FDR < 0.05) among DEGs across TFAM^KD-MEL^cGAS^WT^ versus TFAM^WT-MEL^cGAS^WT^, TFAM^KD-MEL^cGAS^KO^ versus TFAM^KD-MEL^cGAS^WT,^ and TFAM^KD-MEL^cGAS^KO^ versus TFAM^WT-MEL^cGAS^KO^ comparisons; dot color indicates signed -log10(FDR) and dot size reflects gene count. k, Immunoblot analysis of ZBP1, VIPERIN, IFIT3, IL-1β, and STAT1 protein expression across all four genotypes; HSP60 served as loading control.

To determine whether the transcriptional changes observed in TFAM^KD-MEL^ tumors were dependent on cGAS, we performed bulk RNA-seq analysis. cGAS deletion largely reversed the inflammatory gene expression signature associated with Tfam ablation, with TFAM^KD-MEL^cGAS^KO^ tumor transcriptomes shifting toward TFAM^WT-MEL^cGAS^WT^ controls, reflecting broad reversal of the TFAM^KD-MEL^cGAS^WT^ transcriptional state (Fig. 7j). Fold-change comparison demonstrated that genes induced by Tfam ablation and reversed by cGAS deletion were enriched for inflammatory, immunoregulatory, and tissue-remodeling programs (Fig. 7h). Prominent cGAS-dependent genes included inflammatory regulators (*Socs3*, *Rel*, *Nfkbiz*, *Tnfaip3*, *Atf3*, *Runx1*, *Fosb*), immune checkpoint and co-stimulatory molecules (*Pdcd1*, *Ctla4*, *Tnfrsf9*), immune cell-associated genes (*Cd14*, *Cd4*, *Ccr7*, *Rag1*, *Pax5*, *Mzb1*), and extracellular matrix remodeling factors (*Serpine1*, *Ccn1*, *Tgm1*, *Tns4*, *Grem2*) (Fig. 7h). Gene set enrichment analysis confirmed suppression of TNFα/NF-κB signaling, inflammatory response, and hypoxia pathways in TFAM^KD-MEL^cGAS^KO^ tumors (Fig. 7i). Immunoblot analysis validated these findings at the protein level, demonstrating normalization of IL-1β, IFIT3, and phospho-STAT1 toward TFAM^WT-MEL^ levels upon cGAS deletion (Fig. 7j). Collectively, these findings establish that cGAS-mediated innate immune signaling is required for both the accelerated tumor growth and transcriptional remodeling driven by tumor cell–intrinsic mtDNA instability.

## Discussion

Beyond cell-intrinsic metabolic rewiring and regulation of cell growth, mitochondria serve as signaling hubs that regulate cellular homeostasis and, under cellular stress, release mitochondrial DAMPs that can trigger innate immune activation ^3,32,33^. Here, using an autochthonous Braf^V600E^/Pten^-/-^ melanoma model, we demonstrate that tumor cell–intrinsic mtDNA instability drives temporal remodeling of the TME and promotes melanoma progression and dissemination recapitulating features of high-risk human melanoma ^19^. These findings provide a mechanistic framework for how mtDNA instability contributes to tumor progression and help reconcile the apparently opposing, context-dependent effects of mtDNA alterations reported in melanoma ^9–11^.

By inducing melanocyte-specific TFAM ablation in young animals, we modeled tumor cell-intrinsic mtDNA instability while minimizing confounding from systemic aging-associated processes and preserving the native tissue environment and tumor-immune co-evolution that transplantation models preclude. Unlike models involving severe mtDNA perturbation that may predominantly increase metabolic vulnerability, heterozygous TFAM ablation induced mtDNA instability while preserving mitochondrial respiratory function and engaging cGAS-dependent innate immune signaling ^3^, providing an opportunity to examine the consequences of chronic mtDNA stress during tumor progression. Genetic ablation of cGAS reversed both accelerated tumor progression and the associated transcriptomic changes of the TFAM^KD-MEL^ TME, establishing cGAS-dependent innate immune signaling as the central mechanism linking tumor cell-intrinsic mtDNA instability to melanoma progression. Notably, while early TFAM^KD-MEL^ malignant cells exhibited metabolic stress signatures, late-stage tumors showed relative enrichment of OXPHOS and proliferative programs, suggesting adaptation to persistent mtDNA stress during tumor progression. This observation raises the possibility that the extent of mtDNA alteration influences its biological consequences. Severe mtDNA alterations, as seen in mitochondrial disease, may impose metabolic stress and increase tumor-cell immunogenicity, potentially favoring antitumor immune responses. By contrast, mtDNA instability arising from processes such as UV exposure and aging may primarily engage cell-intrinsic stress signaling and reshape the tumor immune microenvironment without causing substantial OXPHOS dysfunction.

We show that mtDNA instability triggered a robust type I IFN response early in tumor development that diminished as tumors progressed. Early palpable TFAM^KD-MEL^ lesions and sgTFAM YUMM1.7 cells exhibited robust type I IFN signaling, whereas malignant cells from established TFAMKD-MEL tumors showed limited cGAS-STING activity and were instead enriched for proliferative and invasive programs. This reframes mtDNA instability not as a static molecular feature but as a dynamic modulator of innate immune signaling that rewires the tumor ecosystem. Despite the antiproliferative activity ^34^, chronic type I IFN signaling has been shown to drive cancer cell stemness and invasive programs^20,21^, raising the possibility that early mtDNA instability-driven IFN signaling contributes to the subsequent acquisition of these programs. The early induction of *Socs3* and *Usp18*, canonical negative-feedback regulators of IFNAR–JAK/STAT1 signaling ^35^, may contribute to cell-intrinsic attenuation of the type I IFN response. Epigenetic silencing of *cGAS* and *STING* has been reported in melanoma cells and may limit sustained innate immune activation in established tumors ^36,37^. Beyond these cell-intrinsic mechanisms, the late-stage dominance of IL-6/JAK/STAT3- and NF-κB-driven programs in the TFAM^KD-MEL^ TME may reflect a broader rewiring of innate immune signaling. A shift from type I IFN signaling toward NF-κB- and JAK/STAT3-associated inflammatory programs downstream of chronic cGAS-STING activation has been reported in the context of chromosomal instability ^13,14^, while IL-6/JAK2/STAT3 signaling has been shown to suppress cGAS-STING-mediated type I IFN expression ^38^. These observations support a model in which cGAS-dependent signaling contributes to immune remodeling of the TFAM^KD-MEL^ TME, while IL-6/JAK2/STAT3 signaling may contribute to attenuation of the type I IFN response during tumor progression. The mechanisms underlying this stage-dependent immune rewiring remain to be defined.

The temporal nature of this immune response has direct clinical implications. The stage-dependent inflammatory macrophage expansion and reprogramming toward immunoregulatory states observed in TFAM^KD-MEL^ tumors parallels aging-associated macrophage dynamics in melanoma, in which inflammatory *Trem1*-high populations give way to *Trem2*-associated immunoregulatory states ^39^, suggesting that TFAM ablation models aspects of mitochondrial aging-associated stress that may contribute to immune remodeling during melanoma progression. Critically, the mtDNA instability-driven early inflammatory macrophage signatures mapped to human melanoma macrophage states linked to favorable survival and response to therapy, whereas the late tissue-adaptive macrophage signatures mapped to states associated with poor prognosis and therapy resistance across independent cohorts. Although anti-PD-1 treatment did not fully abrogate tumor growth, it delayed TFAM^KD-MEL^ tumor progression, supporting a potential susceptibility window associated with the early IFN-rich immune state. The enrichment of efferocytosis-associated macrophage programs at late stage is particularly relevant in this context, as efferocytosis can promote immune tolerance and has been linked to melanoma progression and immunotherapy resistance ^25,30^. Consistent with this temporal model, chronic tumor cell-intrinsic IFN signaling at baseline has been associated with transient ICB response in metastatic melanoma ^40^, and our findings provide potential in vivo mechanistic context for this clinical pattern. Our data further suggests that static measurements of cGAS-STING activity or ISG expression may have limited predictive value because mtDNA-driven immune activation evolves through distinct temporal states with fundamentally different therapeutic implications.

In conclusion, we demonstrate that tumor cell-intrinsic mtDNA instability drives melanoma progression through cGAS-dependent, temporally evolving immune and stromal remodeling. These findings establish mtDNA instability as a dynamic regulator of tumor immune ecosystem whose consequences evolve with disease stage and provide a foundation for prospective validation of immune-stromal signatures as biomarkers of disease progression and treatment response.

## Methods

### Mouse models and husbandry

Tyr::CreERT2; Braf^CA^; Pten*^fl/fl^* mice were obtained from the Bosenberg laboratory (Yale School of Medicine). cGAS*^-/-^* mice (Cgas*tm1d*; JAX strain 026554) and Tfam conditional mice (Tfam*^fl/fl^*; JAX strain 026123) were obtained from The Jackson Laboratory. To generate melanocyte-specific TFAM haploinsufficient melanoma mice, Tfam*^fl/+^*mice were crossed with Tyr::CreERT2; Braf^CA^; Pten*^fl/fl^*mice to produce Braf*^V600E/+^*; Pten*^-/-^*; Tfam*^fl/+^* (TFAM^KD-MEL^) and littermate Braf*^V600E/+^*; Pten*^-/-^*; Tfam*^+/+^* (TFAM^WT-MEL^) controls on a C57BL/6J background. TFAM^KD-MEL^ and TFAM^WT-MEL^ mice were backcrossed onto the cGAS*^-/-^*background to generate TFAM^KD-MEL^cGAS^KO^ and TFAM^WT-MEL^cGAS^KO^ mice. Male and female mice were used across all experimental cohorts. All colonies were maintained under specific pathogen-free conditions. All procedures were approved by the Institutional Animal Care and Use Committee (IACUC). Genotyping was performed using standard PCR protocols.

### Melanoma induction and tumor monitoring

Melanomas were induced in 3-week-old mice by topical application of 4-hydroxytamoxifen (4-OHT; 70% Z-isomer, Sigma) as previously described ^18^. Briefly, mice were anesthetized with isoflurane one day prior to 4-OHT administration, the lower dorsal skin was shaved, and residual hair was removed with a topical depilatory cream (Nair). The following day, 1 µl of 20 mM 4-OHT dissolved in 100% ethanol was applied topically to the exposed skin. Four weeks post induction, dorsal hair was removed using electric trimmers to facilitate tumor monitoring. Tumor growth was measured twice weekly using digital calipers and tumor volume was calculated using the ellipsoid formula (V = W × L × H × 0.5236). Humane endpoints were defined as tumor volume exceeding 2 cm³, tumor ulceration, or signs of systemic distress including impaired ambulation, consistent with IACUC-approved protocols. Survival was defined as the time from palpable tumor formation to humane endpoint, and survival differences between groups were assessed using Kaplan-Meier analysis with log-rank testing.

### Cell lines and lentiviral CRISPR-Cas9-mediated TFAM knockdown

YUMM1.7 murine melanoma cells (Braf^V600E/+^; Pten^-/-^; Cdkn2a^-/-^) and HEK293FT cells were obtained from ATCC and maintained in DMEM supplemented with 10% FBS. For lentiviral production, HEK293FT cells were co-transfected with pLentiCRISPR v2 vectors encoding TFAM-targeting or non-targeting control sgRNAs together with packaging plasmids psPAX2 and pMD2.G using polyethylenimine (PEI) transfection reagent. Viral supernatants were collected at 48 and 72 hours post-transfection, centrifuged at 500 × g for 5 minutes, filtered through a 0.45 µm membrane, and used to transduce YUMM1.7 cells. Stable knockdown lines were selected with puromycin. Two independent sgRNAs targeting *Tfam* were used: sg1TFAM (5′-GGAGCGTGCTAAAAGCACTG-3′) and sg3TFAM (5′-AAAAGCACTGGGGCGCACGG-3′).

A non-targeting sgRNA (sgCTL; 5′-GAAGTTCGAGGGCGACACCC-3′) served as a control. TFAM knockdown was confirmed by immunoblotting and immunofluorescence as described below.

### Immunofluorescence and multiplex immunohistochemistry

Cells were seeded on glass coverslips in 24-well plates and cultured overnight to 50-70% confluency. Cells were washed with PBS, fixed in 4% paraformaldehyde for 20 minutes, permeabilized with 0.1% Triton X-100 in PBS for 5 minutes at room temperature, and blocked with 5% FBS in PBS for 1 hour. Coverslips were incubated with primary antibodies: anti-dsDNA (MilliporeSigma, AC-30-10; 1:300), anti-HSP60 (Proteitech, 2F10E7; 1:300), and anti-TFAM (MilliporeSigma, ABE483; 1:800) for 1 hour at room temperature. After three washes in PBS, coverslips were incubated with species-appropriate fluorescent secondary antibodies, Rhodamine Red-X anti-mouse IgM (1:600), Alexa Fluor 488 anti-rabbit (1:600), and Alexa Fluor 647 anti-goat (1:600), for 1 hour at room temperature in the dark. Coverslips were mounted using antifade mounting medium with DAPI (Invitrogen) and imaged on Nikon fluorescence microscope with a 60× oil immersion objective using NIS-Elements.

### Mitochondrial DNA copy number analysis

Total DNA was isolated from snap-frozen tissues or cell pellets using the Quick-DNA Microprep Kit (Zymo Research, R1051) according to the manufacturer’s instructions. DNA concentration was quantified using a NanoDrop spectrophotometer, and approximately 10 ng of template DNA was used per qPCR reaction performed using PerfeCTa SYBR Green FastMix on a QuantStudio real-time PCR system. Relative mtDNA copy number was determined by amplifying the mitochondrial genes and normalized to the nuclear-encoded reference gene *Tert*. Relative mtDNA abundance was calculated using the 2 t method, where ΔCt = Ct(*mt-Nd1*) – Ct (*Tert*).

### Bulk mRNA sequencing

Snap-frozen tumor samples were pulverized to a fine powder using pre-chilled grinders and lysed in TRIzol reagent (Invitrogen). RNA was extracted according to the manufacturer’s instructions. For cell lines, RNA was extracted using the Zymo RNA Miniprep Kit. RNA concentration and quality were assessed using a NanoDrop spectrophotometer (Thermo Scientific) and the RNA ScreenTape Assay (Agilent Technologies). Stranded mRNA libraries were prepared using the KAPA mRNA HyperPrep Kit and quantified using the D5000 ScreenTape Assay and Qubit dsDNA HS Assay. Libraries were sequenced on an Illumina NovaSeq X Plus platform with paired-end 150 bp reads. Reads were aligned to the mm10 mouse reference genome using STAR. Genes with zero counts across all samples were removed. Library size normalization was performed using the trimmed mean of M-values (TMM) method, and expression values were transformed to log counts per million (CPM + 1) for downstream analysis. Differential expression analysis was performed using the DESeq2 package in R (v 1.46.0), with pairwise comparisons between experimental conditions; genes were considered differentially expressed at an adjusted p-value < 0.05 and |log fold change| > 1 unless otherwise stated. Gene Set Enrichment Analysis (GSEA) was performed on genes ranked by log□ fold change using mouse Hallmark gene sets from MSigDB with the fgsea R package. KEGG and Reactome pathway analysis was performed using the clusterProfiler package.

### Single-cell RNA sequencing and analysis

Whole tumors from early-stage melanomas (300-500 mm³; 42 days post-induction) and representative sections of late-stage melanomas (1,000-1,500 mm³; 55-60 days post-induction) were mechanically minced and enzymatically dissociated using the Mouse Tumor Dissociation Kit and gentleMACS Octo Dissociator (Miltenyi Biotec) according to the manufacturer’s 37C_m_TDK_1 protocol. Cell suspensions were filtered through 70-μm strainers and subjected to red blood cell lysis (Thermo Fisher Scientific). Dead cells and debris were removed using Dead Cell and Debris Removal Kits (Miltenyi Biotec). Cells were resuspended in PBS containing 0.04% BSA, assessed for viability, and single-cell RNA-seq libraries were prepared using the Chromium Single Cell 3′ Reagent Kit v3.1 (10x Genomics) and sequenced on an Illumina HiSeq X10 platform at the Jackson Laboratory Single Cell Biology Core Facility. Raw sequencing data were processed using Cell Ranger and aligned to the mm10 mouse reference genome to generate filtered gene–barcode matrices. Quality control and downstream analyses were performed using Seurat (v5.2.0). Cells were filtered based on the number of detected genes (<300 or >9,000), UMI counts (<800 or >70,000), or mitochondrial gene content (>15%), resulting in 152,954 high-quality cells across ten samples (mean: 15,295 per sample; range: 10,388–17,983; retention rate: 92.2%).

Gene expression was normalized using SCTransform (v2), with mitochondrial gene content and cell-cycle scores regressed during normalization. Batch effects were corrected using Harmony on the top 50 principal components. Doublets and low-quality clusters were identified using high-resolution overclustering (resolution = 2.0) and removed based on lineage-inconsistent marker co-expression (e.g., *Col1a1* with *Ptprc,* or *Cd3d* with *Cd19*) and low transcriptomic complexity. The filtered dataset was re-clustered using the first 40 principal components at a resolution of 0.3, yielding 23 transcriptionally distinct populations, which were annotated into major cell types including melanoma, fibroblast, endothelial, and immune compartments based on canonical markers. Major cell types including melanoma cells, CAFs, myeloid cells, and T/NK cells were independently subsetted for lineage-specific re-clustering and refined annotation. Differential gene expression analysis was performed on the SCT assay using Seurat’s FindMarkers function with the MAST-based test. Differentially expressed genes were defined using an absolute log-fold change > 0.25 and Benjamini–Hochberg-adjusted *P* < 0.05.

Pathway enrichment analysis was performed using fgsea and clusterProfiler against MSigDB Hallmark, KEGG, Gene Ontology Biological Process, and Reactome gene sets. For pseudobulk pathway enrichment, single-cell counts were aggregated per sample within each cell type, differential expression between genotypes was tested using edgeR with default parameters, and genes were ranked by a signed statistic combining fold-change direction and statistical significance for fgsea. Cell-level gene signature and pathway activities were quantified using UCell with default parameters, and group differences were assessed using two-sided Wilcoxon rank-sum tests with Benjamini-Hochberg correction. MDSC gene signatures were derived from a previously published study ^24^ and scored using UCell on the SCT assay. CAF subcluster pathway activity was quantified using UCell based on MSigDB Hallmark gene sets. Mean pathway scores were computed per CAF subcluster, z-score normalized, and used to construct a subcluster-by-subcluster Pearson correlation matrix to assess functional similarity; hierarchical clustering was performed using Ward.D2 linkage.

Cell-cell communication was inferred using CellChat (v2) applied to normalized gene expression data across all major cell populations. Differential interaction strength and pathway-level information flow between TFAM^KD-MEL^ and TFAM^WT-MEL^ tumors were computed using the built-in differential analysis framework. Macrophage developmental trajectories were reconstructed using Monocle3 (v1.3.1), with root cells defined as those in the top decile of *Ly6c2, Vcan, Sell* expression corresponding to classical monocytes. Gene expression dynamics along pseudotime were visualized using LOESS smoothing stratified by genotype.

### Macrophage signature scoring and clinical correlation analysis

Differentially expressed genes between TFAM^KD-MEL^ and TFAM^WT-MEL^ macrophages were identified using MAST on the SCT assay, with sex included as a latent variable. Differentially expressed genes were ranked by log FC and mapped to human orthologs using the homologene package (v1.4.68). The top 100 upregulated genes at each time point were retained to define the early and late TFAM^KD-MEL^ macrophage signatures. Processed scRNA-seq data from the Yang et al. melanoma ICI meta-analysis cohort ^31^ were obtained and subsetted to monocyte/macrophage populations based on the original cell-type annotations. Cells were re-normalized and re-clustered using a standard Seurat v5 workflow, with patient-level batch correction performed using Harmony. Clusters were annotated based on canonical marker genes identified by differential expression, yielding six myeloid subtypes as described in the Results. TFAM^KD-MEL^ macrophage signature scores were calculated for individual cells using UCell. For each signature, scores in MRC1□ Mac were compared with those across all other myeloid subtypes combined using two-sided Wilcoxon rank-sum tests. MRC1□ Mac signature scores were additionally compared between ICI responders and non-responders using two-sided Wilcoxon rank-sum tests. For each of the six myeloid subtypes, frequency was calculated as the percentage of total myeloid cells within each sample and compared between ICI responders and non-responders using an exact two-sided Wilcoxon rank-sum test implemented in the coin R package. P values across the six subtype comparisons were adjusted for multiple testing using the Benjamini-Hochberg (FDR) method. This analysis was restricted to samples receiving ICI-containing treatment regimens, including anti-PD-1 monotherapy, anti-PD-1/anti-CTLA-4 combination therapy, and other combination regimens containing ICI. Treatment response was defined according to the response annotation provided for each sample.

scRNA-seq data from melanoma patients treated with BRAF/MEK-targeted therapy were obtained from Vasilevska et al. (GSE229908). Macrophages annotated as therapy-sensitive (TTS) or therapy-resistant (TTR) were subsetted and scored independently for the early and late TFAM-^KD-MEL^ macrophage signatures using UCell (v2.4). Signature scores were compared between TTS and TTR macrophages using two-sided Wilcoxon rank-sum tests.For each time point, upregulated (KD-like) and downregulated (WT-like) macrophage genes were selected independently. Signature activity was quantified in the TCGA-SKCM bulk RNA-seq cohort (n = 460; TCGA Research Network), comprising primary and metastatic melanomas, using single-sample gene set enrichment analysis (ssGSEA) implemented in GSVA (v1.48), with score normalization enabled. A composite macrophage activity score was calculated for each patient as the difference between KD-like and WT-like ssGSEA enrichment scores. Patients were independently stratified at the cohort median into high-score (KD-like) and low-score (WT-like) groups for the early and late signatures. Overall survival was assessed using Kaplan–Meier analysis with a two-sided log-rank test. Cox proportional hazards regression using the composite score as a continuous variable was performed as a sensitivity analysis.

### Flow cytometry

Tumors were harvested, finely minced with sterile scissors, and enzymatically dissociated using the gentleMACS Octo Dissociator (Miltenyi Biotec) with the 37C_m_TDK_1 protocol. Single cell suspensions were filtered through a 70 µm strainer, incubated with LIVE/DEAD Fixable Blue Dead Cell Stain (Invitrogen) and anti-mouse CD16/CD32 Fc-blocking antibody (2.4G2) in PBS for 20 minutes at 4°C, then stained with fluorochrome-conjugated antibodies in FACS buffer (2% FBS, 2 mM EDTA in PBS) for 30 minutes at 4°C. The following antibodies were used for surface staining: CD3-PE-CF594 (17A2), CD4-BV805 (RM4-4), CD8-BV510 (53-5.8), NK1.1-Spark NIR 685 (S17016D), CD25-BV785 (3C7), CD19-BB700 (6D5), PD-1-PE (29F.1A12), KLRG1-PE-Fire700 (2F1/KLRG1), F4/80-PE-Cy7 (BM8), MHC II-BV711 (AF6-120.1), CD11c-FITC (N418), Ly6G-PE (1A8), Ly6C-BV650 (HK1.4), CD206-PE-Cy5 (C068C2), and PD-L1-BV421 (10F.9G2) from BioLegend; CD45-BUV737 (30-F11), CD86-BV480, CXCR2-PE-CF594 (V48-2310), and CTLA-4-PE-Cy7 from BD Biosciences; and CD11b-APC-Cy7 (M1/70) from Tonbo Biosciences. For FOXP3 intracellular staining, cells were fixed and permeabilized using the eBioscience Foxp3 Transcription Factor Staining Buffer Set (00-5523-00, Thermo Fisher Scientific) for 30 minutes at room temperature, followed by staining with Foxp3-AF647 (MF-14, BioLegend) in permeabilization buffer. Data were acquired on a BD FACSymphony A5 SE spectral flow cytometer and analyzed using FlowJo (v10.10).

### Immunoblotting

Snap-frozen tumor samples were ground to a fine powder using pre-chilled grinders and lysed in NP-40 lysis buffer supplemented with protease inhibitors (Thermo Fisher Scientific). Total protein concentration was determined using the Micro BCA Protein Assay Kit (Thermo Fisher Scientific). Equal amounts of protein (15-30 µg) were resolved on 10-12% SDS-PAGE gels and transferred to PVDF membranes (Bio-Rad). Membranes were air-dried to return to a hydrophobic state and incubated overnight at 4°C with the following primary antibodies: TFAM (ABE483, MilliporeSigma), cGAS (D3080, CST), STING (D2P2F, CST), TBK1 (D1B4, CST), phospho-TBK1 Ser172 (D52C2, CST), IRF3 (D83B9, CST), NF-κB p65 (D14E12, CST), ZBP1 (aa 1–411, AdipoGen), IFIT3 (gift from G. Sen, Cleveland Clinic), IL-1β (3A6, CST), Gasdermin D (96458, CST), Caspase-1 p20 (AdipoGen), MTCO1 (1D6E1A8, Abcam), OXPHOS Rodent Cocktail (ab110413, Abcam), and HSP60 (2F10E7, Proteitech). After three washes in PBS, membranes were incubated with HRP-conjugated secondary antibodies for 1 hour at room temperature. Chemiluminescent signal was developed using Immobilon Crescendo HRP Substrate and detected by chemiluminescence imaging.

### In vivo PD-1 blockade

Mice bearing established tumors (50-100 mm³) were stratified by tumor volume and randomized into treatment groups. Mice received 200 µg of anti-mouse PD-1 antibody (29F.1A12, Bio X Cell) or PBS control intraperitoneally in 200 µl sterile PBS twice weekly for the duration of the experiment. Tumor growths were measured twice weekly using digital calipers and volume calculated as described above. Mice were euthanized upon reaching humane endpoints and survival was assessed using Kaplan–Meier analysis with log-rank testing.

### Statistical analysis

Unless otherwise specified, statistical analyses of tumor growth, flow cytometry, and other non-transcriptomic data were performed using GraphPad Prism (v10). Comparisons between two groups were conducted using two-tailed unpaired Student’s t-tests. For comparisons among three or more groups, one-way ANOVA followed by Tukey’s multiple comparisons test was applied. Tumor growth curves were compared using two-way repeated-measures ANOVA with Sidak’s multiple comparisons test. Survival analyses were performed using the Kaplan-Meier method with log-rank testing. All data are presented as mean ± standard error of the mean (SEM). A p-value < 0.05 was considered statistically significant.

## Data and code availability

Raw bulk RNA-sequencing data generated in this study have been deposited in the NCBI Gene Expression Omnibus (GEO) under accession number GSE[XXXXXX]. Raw and processed single-cell RNA-sequencing data have been deposited in GEO under accession number GSE[XXXXXX]. Publicly available datasets used in this study include human melanoma scRNA-seq data from Yang et al. ^31^, Sade-Feldman et al. (GSE120575), Jerby-Arnon et al. (GSE115978), and Vasilevska et al. (Zenodo: https://zenodo.org/records/11208953). TCGA-SKCM RNA-seq data were obtained from the Genomic Data Commons portal using the TCGAbiolinks R package (v2.38.0). All analysis code used to generate the results and figures in this study is available from the corresponding author upon reasonable request.

## Acknowledgements

We gratefully acknowledge support from the Jackson Laboratory Scientific Services, including the Single Cell Biology Laboratory, Genome Technologies, Protein Sciences, Histopathology Sciences, Microscopy, and Computational Sciences.

## Funding

This work was supported by the Cancer Prevention and Research Institute of Texas (CPRIT) under grant number RP170734. Additional support was provided by the National Institutes of Health under Award Numbers R01HL148153 and P30CA034196. The content is solely the responsibility of the authors and does not necessarily represent the official views of the National Institutes of Health. A.A.B. was supported by a Brooks Scholar Award from The Jackson Laboratory Cancer Center. A.S. is supported by a JAX Scholar Award and American Cancer Society Postdoctoral Fellowship PF-26-1594975-01. J.F.B. was supported by American Cancer Society Postdoctoral Fellowship PF-25-1420432-01.

## Author contributions

Investigation: A.A.B., L.C.W., A.D.W., Y.L., L.F.A., D.G.T.V., P.S.G., M.P., and A.P.W.; Methodology: A.A.B., L.C.W., A.D.W., Y.L., A.S., J.F.B., J.C.R., G.S.S., M.B., and A.P.W.; Visualization: A.A.B., L.C.W., A.D.W., and A.P.W.; Writing,: A.A.B. and A.P.W.; Funding acquisition: A.A.B., L.C.W., and A.P.W. All authors read and approved the manuscript.

## Extended Data Figures

**Extended Data Fig. 1.**
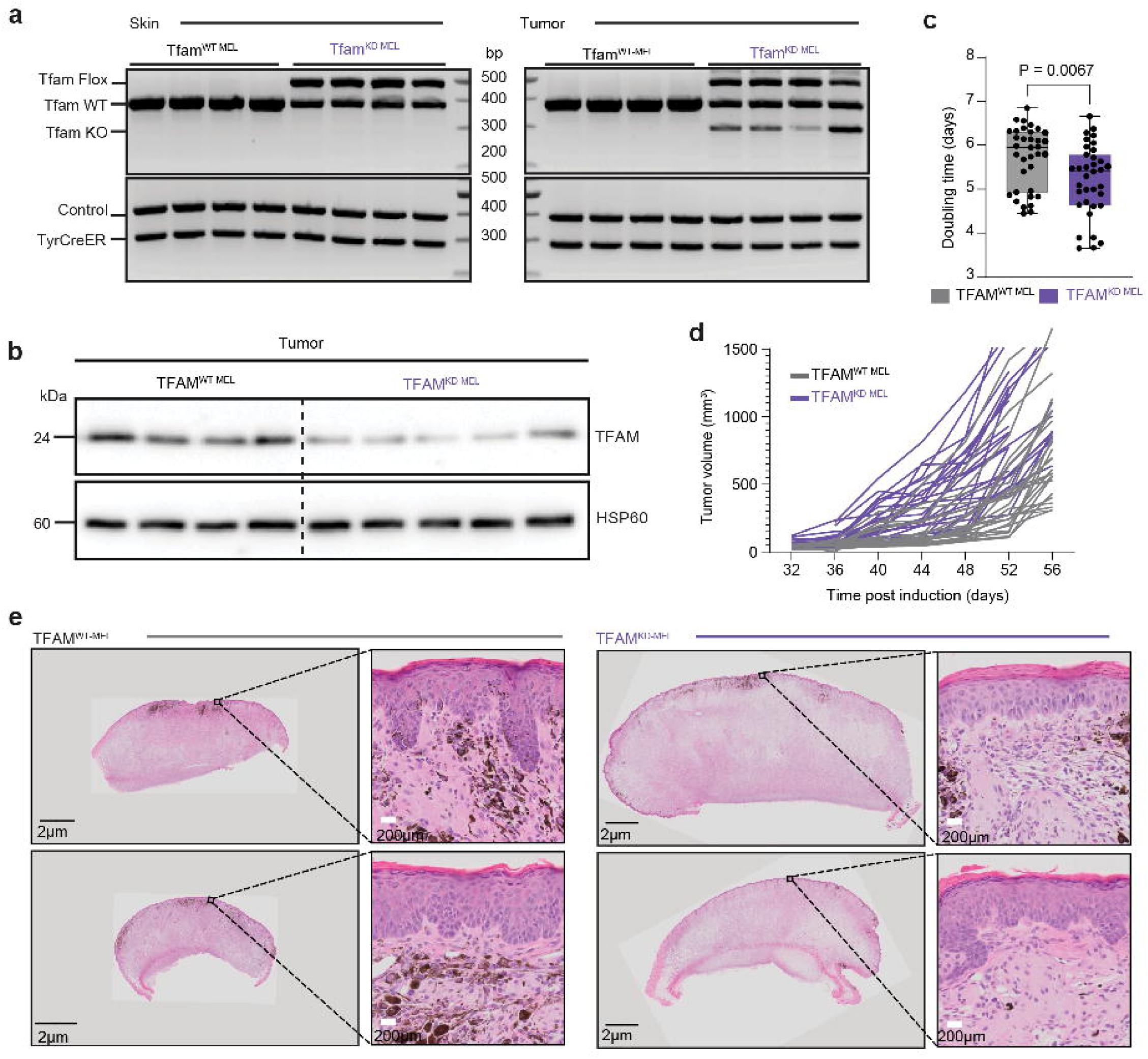
Validation of Tfam knockdown and tumor growth kinetics in the TFAM^KD-MEL^ melanoma model. a, PCR-based genotyping of *Tfam* in skin and tumor tissues. Bands correspond to floxed (Flox), wild-type (WT), and recombined knockout (KO) *Tfam* alleles. b, Immunoblot analysis of TFAM protein expression in early-stage tumors; HSP60 was used as a loading control. c, Tumor volume doubling time following palpable tumor detection (n = 50-60 mice per group). d, Representative individual tumor growth curves. e, Representative hematoxylin and eosin (H&E)-stained tumor sections showing melanoma morphology at low and high magnification. Scale bars, 200 μm and 50 μm.

**Extended Data Fig. 2.**
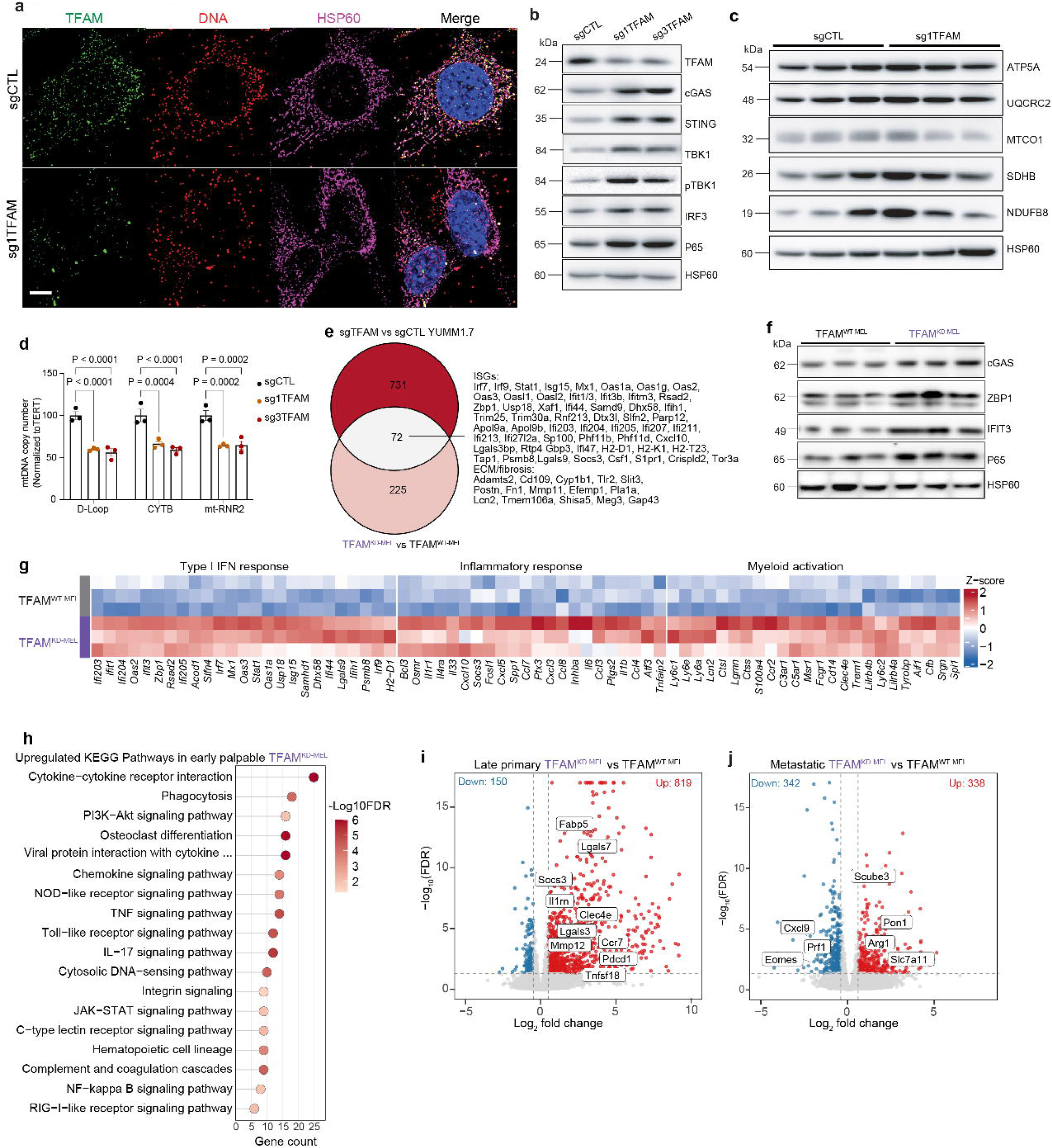
TFAM knockdown induces mtDNA instability and activates cGAS-STING-mediated type I IFN signaling in melanoma. a, Representative immunofluorescence images of TFAM (green), DNA (red) and HSP60 (magenta) in sgTFAM and sgCTL YUMM1.7 cells. b,c, Immunoblots of cGAS-STING signaling components (b) and OXPHOS complex subunits (c); HSP60 served as a loading control. d, Relative mtDNA copy number measured at *D-loop*, *CYTB* and *mt-RNA2*, normalized to the nuclear gene *Tert*. e, Venn diagram showing the overlap between DEGs identified in early palpable TFAM^KD-MEL^ tumors and sgTFAM YUMM1.7 cells relative to their respective controls. f, Immunoblots of cGAS, TBK1, ZBP1, IFIT3, VIPERIN and p65 protein in early TFAM^KD-MEL^ and TFAM^WT-MEL^ tumors; HSP60 served as a loading control. g, Heatmap of DEGs in early palpable TFAM^KD-MEL^ versus TFAM^WT-MEL^ tumors (FDR < 0.05, |log□ FC| > 0.6), highlighting genes associated with type I IFN signaling and myeloid activation. h, KEGG pathways significantly enriched among DEGs in early palpable TFAM^KD-MEL^ tumors relative to TFAM^WT-MEL^ tumors; dot size represents gene set size and color indicates NES. i,j, Volcano plots showing DEGs between TFAM^KD-MEL^ and TFAM^WT-MEL^ primary (i) and metastatic (j) tumors collected at experimental endpoint. Genes upregulated in TFAM^KD-MEL^ tumors (FDR < 0.05, log□ FC > 1) are shown in red, and genes enriched in TFAM^WT-MEL^ tumors are shown in blue; representative inflammatory and immunoregulatory genes (i) and invasion-associated and immune-effector genes (j) are highlighted.

**Extended Data Fig. 3.**
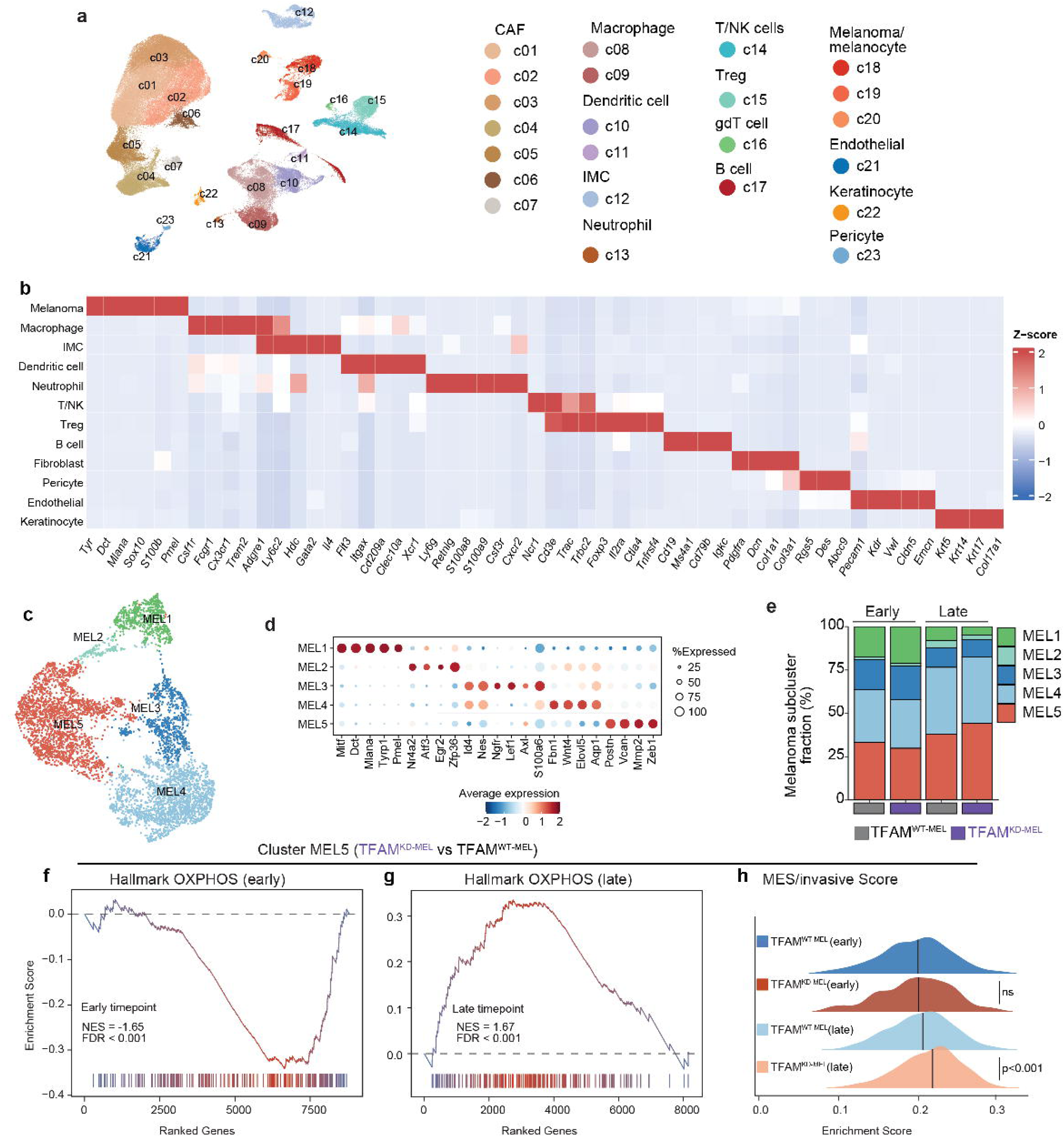
**Temporal metabolic and transcriptional state alterations in TFAM knockdown melanoma cells**. a, UMAP plot of the integrated scRNA-seq dataset from early and late TFAM^KD-MEL^ and TFAM^WT-MEL^ tumors, colored by clusters. b, Heatmap showing marker gene expressions used to annotate major cell populations. Color indicates scaled mean expression (z-score). c, UMAP visualization of Sox10 melanoma cells from TFAM^WT-MEL^ and TFAM^KD-MEL^ tumors. d, Dot plot showing marker gene expression across melanoma subclusters (MEL1-MEL5). Dot size indicates the fraction of expressing cells and color represents scaled mean expression (z-score). e, Stacked bar chart showing the relative abundance of melanoma subclusters in TFAM^WT-MEL^ and TFAM^KD-MEL^ tumors at early and late time points. f, g, GSEA plots showing enrichment of the Hallmark Oxidative Phosphorylation gene set in MEL5 cells at the early (NES = -1.65, FDR < 0.001) (f) and late (NES = 1.67, FDR < 0.001) (g) time points. h, Ridge plot showing MES/invasive gene signature scores in MEL5 cells from TFAM^WT-MEL^ and TFAM^KD-MEL^ tumors at early and late time points. Statistical significance was assessed using a two-sided Wilcoxon rank-sum test (P < 0.001; ns, not significant).

**Extended Data Fig. 4.**
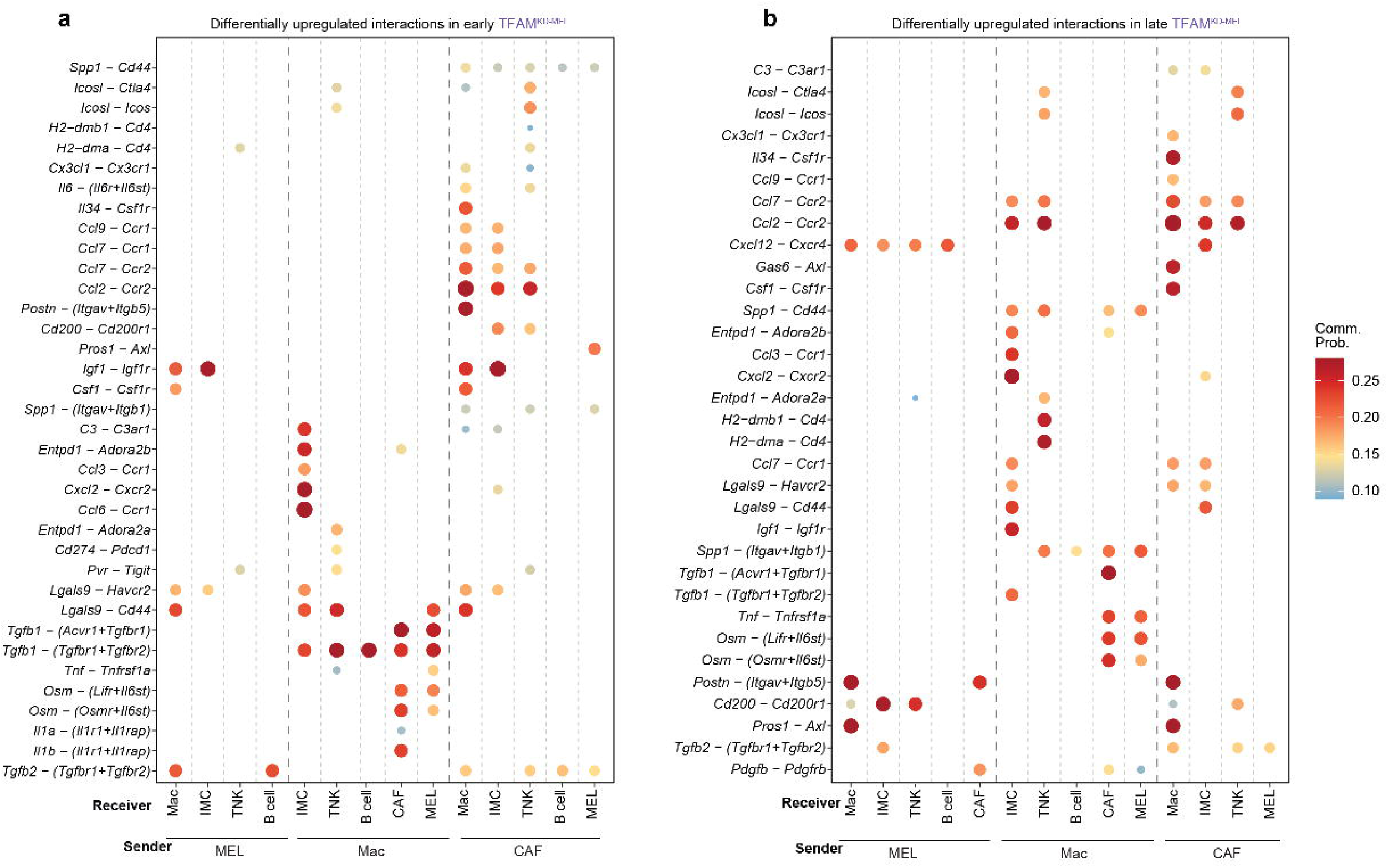
mtDNA instability reshapes intercellular communication in the TME. a,b, Dot plots of ligand-receptor interactions significantly enriched in TFAM^KD-MEL^ tumors at early (a) and late (b) time points across indicated sender-receiver cell pairs. Dot size and color represent communication probability. Interactions are ranked by maximum communication probability across all source cell types; the top 25 interactions are shown, excluding collagen and laminin structural networks.

**Extended Data Fig. 5.**
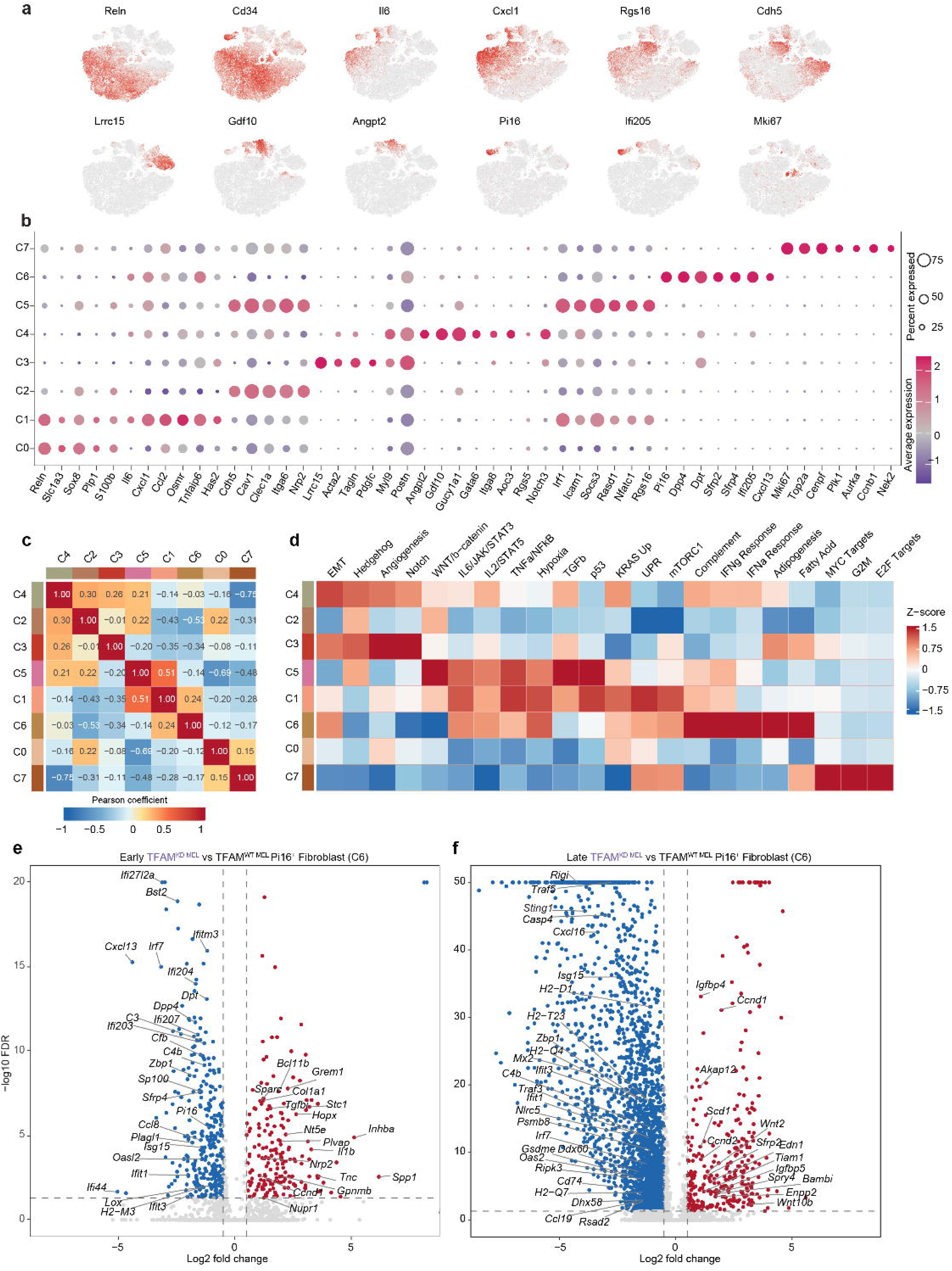
CAF subcluster characterization. a, Feature plots showing representative marker gene expression used to define CAF subclusters. b, Dot plot showing marker gene expression across CAF subclusters. Dot size indicates the fraction of expressing cells and color represents scaled mean expression (z-score). c, Heatmap showing Pearson correlation coefficients across 50 Hallmark pathways among CAF subclusters. Color indicates Pearson correlation from -1 (blue) to 1 (red). Subclusters are ordered by Ward.D2 hierarchical clustering. d, Heatmap showing functional clustering of CAF subclusters based on Hallmark pathway activity scores. Values are column-wise z-scored and capped at ±1.5. e, Volcano plots showing DEGs between TFAM^KD-MEL^ and TFAM^WT-MEL^ tumors within Pi16□ (C6) fibroblasts at early and late time points. The x-axis shows log□ fold change, and the y-axis shows-log□□(adjusted *P* value); selected genes are highlighted.

**Extended Data Fig. 6.**
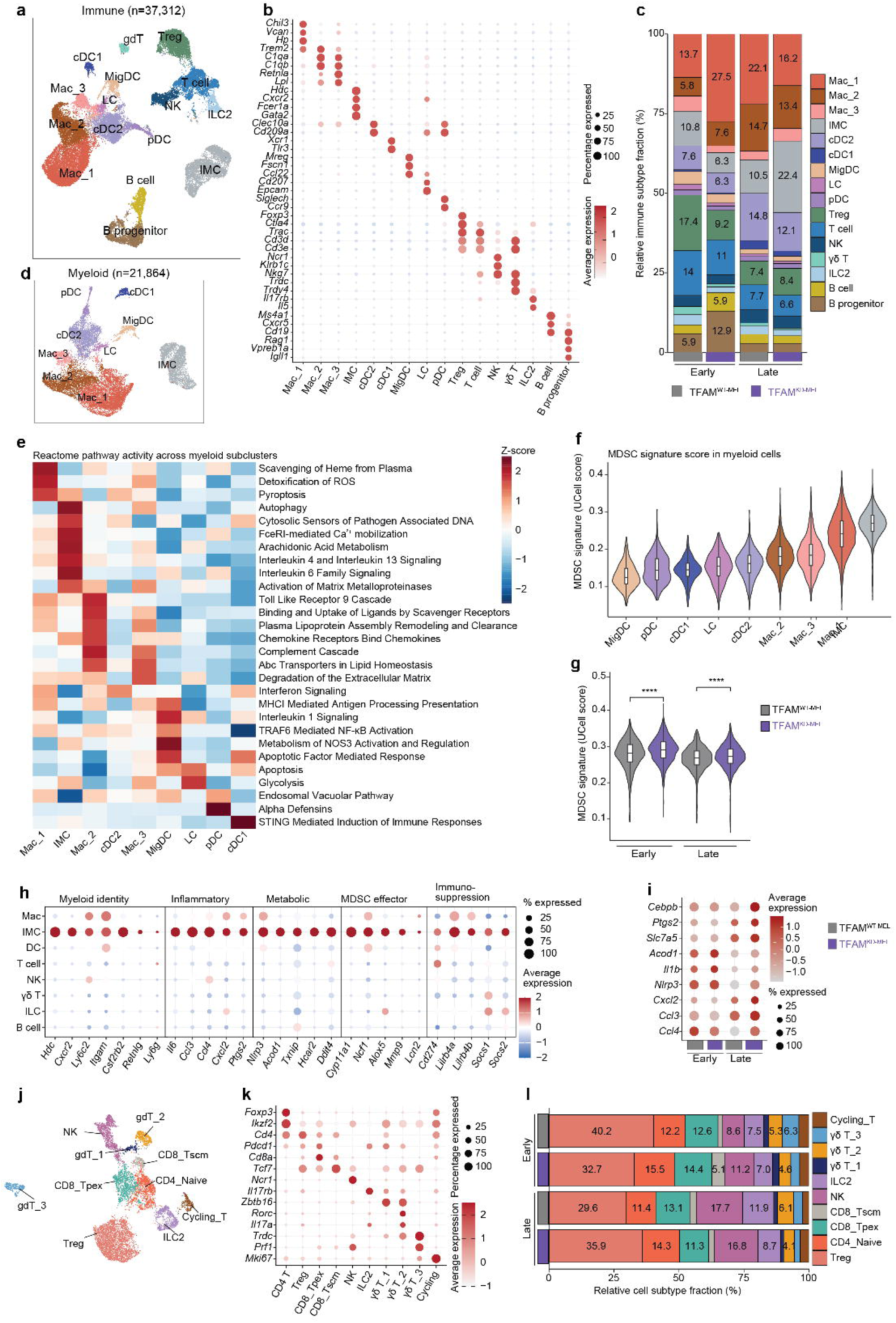
mtDNA instability drives stage-dependent immune remodeling characterized by early Treg reduction and late enrichment of myeloid populations and Tregs. **a**, UMAP of immune cell populations (n = 31,957) from integrated early and late scRNA-seq datasets, comprising macrophages (n = 12,572), five dendritic cell subsets (cDC1, cDC2, mDC, LC, pDC; n = 6,507), IMC (n = 4,834), Treg (n = 3,834), conventional CD4□ and CD8a□ T cells (n = 3,131), NK cells (n = 1,346), γδT (n = 928), ILC (n = 865), and B cells (n = 3,707). b, Dot plot of marker genes enrichment across immune subtypes. c, Bar plots show the relative percentage of immune subtypes at early and late time points. d, UMAP plot of reclustered myeloid cells from a. e, Heatmap displaying the Reactome pathway activities for different myeloid subtypes. f, Violin plots showing MDSC related gene signature ^24^scores across all myeloid subclusters, ranked by median score, with boxplots overlaid to indicate distribution quartiles. g, Violin plots comparing UCell MDSC gene signature scores between TFAM^KD-MEL^ and TFAM^WT-MEL^ IMCs at early and late time points. Statistical significance was assessed using a two-sided Wilcoxon rank-sum test (P < 0.05, **P < 0.01, ***P < 0.001; ns, not significant). h, Dot plot of DEGs in IMCs relative to other major immune cell populations, grouped by functional category. i, Dot plot of selected upregulated genes in IMC cluster from TFAM^KD-MEL^ tumors, including inflammasome components (*Il1b*, *Nlrp3*) and transcriptional regulators associated with myeloid suppressive function (*Cebpb*). j, UMAP of T cells, NK cells, and innate lymphoid cells, colored by cell type. k, Marker gene enrichment across T cells, NK cell, and innate lymphoid cell subtypes. l, Stacked bar plots showing the percentage of each cell type shown in j, normalized to total cells in TFAM^KD-MEL^ and TFAM^WT-MEL^ tumors per time point.

**Extended Data Fig. 7.**
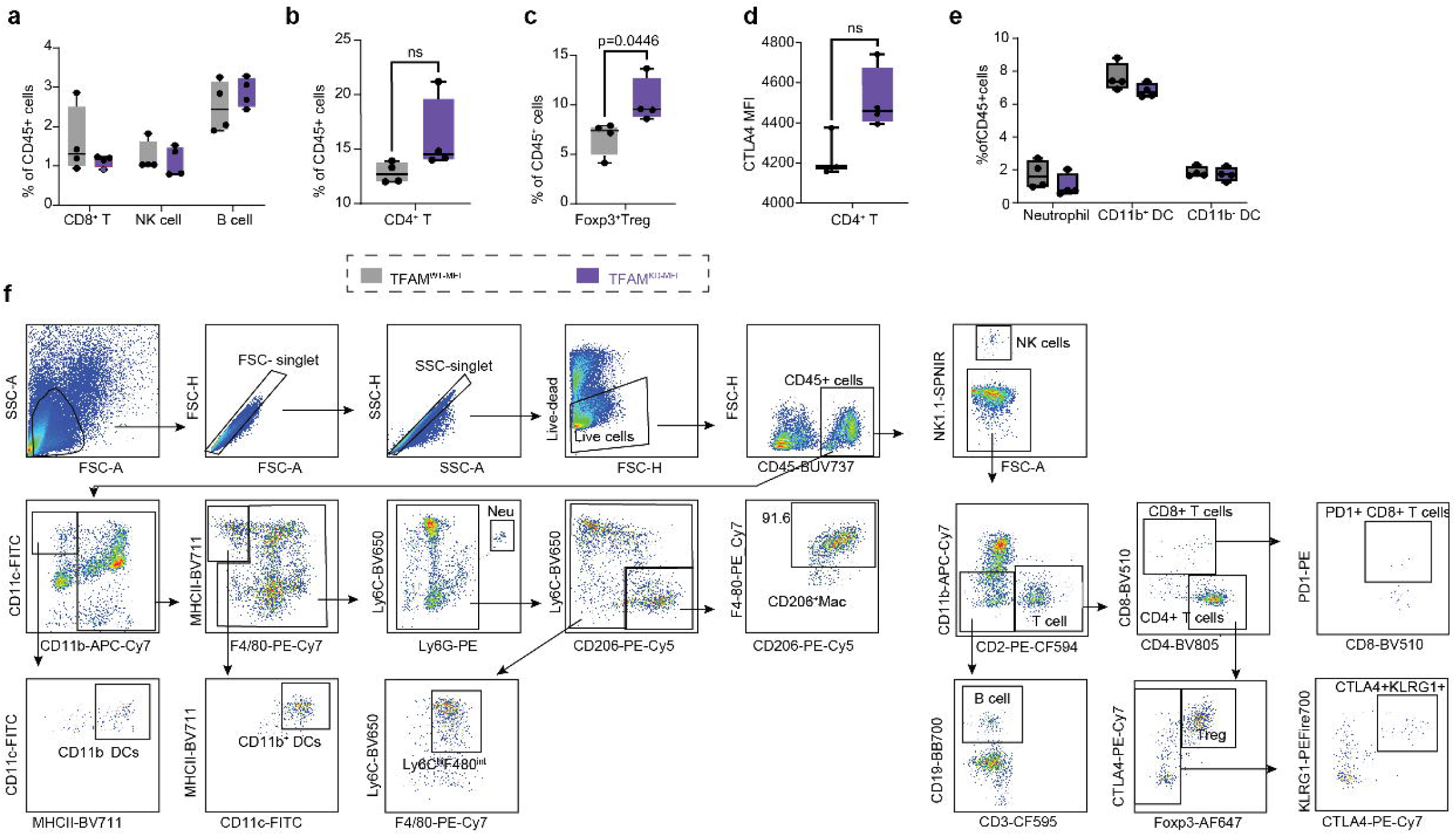
**Flow cytometry demonstrated increased Tregs in TFAM^KD-MEL^ tumors. a-d**, Flow cytometry analysis of immune cell populations in TFAM^KD-MEL^ and TFAM^WT-MEL^ tumors collected at the late time point, depicting increased Tregs in TFAM^KD-MEL^ tumors; unpaired t-test and two-way ANOVA with multiple comparisons correction were used to assess statistical significance (ns, not significant). **e**, Flow cytometry gating strategy for identifying myeloid and lymphoid immune cell populations in tumor tissue.

## Notes

### Competing Interest Statement

The authors have declared no competing interest.

## References

1 Gorelick, A. N. et al. Respiratory complex and tissue lineage drive recurrent mutations in tumour mtDNA. Nature Metabolism 3, 558–570, doi:10.1038/s42255-021-00378-8 (2021).

2 Yuan, Y. et al. Comprehensive molecular characterization of mitochondrial genomes in human cancers. Nature Genetics 52, 342–352, doi:10.1038/s41588-019-0557-x (2020).

3 West, A. P. et al. Mitochondrial DNA stress primes the antiviral innate immune response. Nature 520, 553–557, doi:10.1038/nature14156 (2015).

4 Stewart, J. B. & Chinnery, P. F. Extreme heterogeneity of human mitochondrial DNA from organelles to populations. Nat Rev Genet 22, 106–118, doi:10.1038/s41576-020-00284-x (2021).

5 VanPortfliet, J. J. et al. Caspase-11 drives macrophage hyperinflammation in models of Polg-related mitochondrial disease. Nat Commun 16, 4640, doi:10.1038/s41467-025-59907-8 (2025).

6 Swalwell, H., Latimer, J., Haywood, R. M. & Birch-Machin, M. A. Investigating the role of melanin in UVA/UVB- and hydrogen peroxide-induced cellular and mitochondrial ROS production and mitochondrial DNA damage in human melanoma cells. Free Radic Biol Med 52, 626–634, doi:10.1016/j.freeradbiomed.2011.11.019 (2012).

7 Bowman, A. & Birch-Machin, M. A. Mitochondrial DNA as a Sensitive Biomarker of UV-Induced Cellular Damage in Human Skin. Methods Mol Biol 2277, 345–356, doi:10.1007/978-1-0716-1270-5_21 (2021).

8 Poetsch, M., Dittberner, T., Petersmann, A. & Woenckhaus, C. Mitochondrial DNA instability in malignant melanoma of the skin is mostly restricted to nodular and metastatic stages. Melanoma Res 14, 501–508, doi:10.1097/00008390-200412000-00010 (2004).

9 Ikeda, H. et al. Immune evasion through mitochondrial transfer in the tumour microenvironment. Nature 638, 225–236, doi:10.1038/s41586-024-08439-0 (2025).

10 Monson, K. R. et al. Inherited mitochondrial genetics as a predictor of immune checkpoint inhibition efficacy in melanoma. Nature Medicine 31, 2385–2396, doi:10.1038/s41591-025-03699-3 (2025).

11 Mahmood, M. et al. Mitochondrial DNA mutations drive aerobic glycolysis to enhance checkpoint blockade response in melanoma. Nat Cancer 5, 659–672, doi:10.1038/s43018-023-00721-w (2024).

12 Shelton, S. D. et al. Pathogenic mitochondrial DNA mutations inhibit melanoma metastasis. Sci Adv 10, eadk8801, doi:10.1126/sciadv.adk8801 (2024).

13 Li, J. et al. Non-cell-autonomous cancer progression from chromosomal instability. Nature 620, 1080–1088, doi:10.1038/s41586-023-06464-z (2023).

14 Hong, C. et al. cGAS-STING drives the IL-6-dependent survival of chromosomally instable cancers. Nature 607, 366–373, doi:10.1038/s41586-022-04847-2 (2022).

15 McCastlain, K. et al. Somatic mtDNA mutations at intermediate levels of heteroplasmy are a source of functional heterogeneity among primary leukemic cells. Sci Adv 11, eadt3873, doi:10.1126/sciadv.adt3873 (2025).

16 Araujo, L. F. et al. Mitochondrial transcription factor A (TFAM) shapes metabolic and invasion gene signatures in melanoma. Sci Rep 8, 14190, doi:10.1038/s41598-018-31170-6 (2018).

17 Li, Y. et al. TFAM downregulation promotes autophagy and ESCC survival through mtDNA stress-mediated STING pathway. Oncogene 41, 3735–3746, doi:10.1038/s41388-022-02365-z (2022).

18 Dankort, D. et al. Braf(V600E) cooperates with Pten loss to induce metastatic melanoma. Nat Genet 41, 544–552, doi:10.1038/ng.356 (2009).

19 Gershenwald, J. E. et al. Melanoma staging: Evidence-based changes in the American Joint Committee on Cancer eighth edition cancer staging manual. CA Cancer J Clin 67, 472–492, doi:10.3322/caac.21409 (2017).

20 Musella, M. et al. Type I IFNs promote cancer cell stemness by triggering the epigenetic regulator KDM1B. Nature Immunology 23, 1379–1392, doi:10.1038/s41590-022-01290-3 (2022).

21 Qiu, J. et al. Cancer cells resistant to immune checkpoint blockade acquire interferon-associated epigenetic memory to sustain T cell dysfunction. Nature Cancer 4, 43–61, doi:10.1038/s43018-022-00490-y (2023).

22 Cords, L. et al. Cancer-associated fibroblast classification in single-cell and spatial proteomics data. Nature Communications 14, 4294, doi:10.1038/s41467-023-39762-1 (2023).

23 Forsthuber, A. et al. Cancer-associated fibroblast subtypes modulate the tumor-immune microenvironment and are associated with skin cancer malignancy. Nat Commun 15, 9678, doi:10.1038/s41467-024-53908-9 (2024).

24 Alshetaiwi, H. et al. Defining the emergence of myeloid-derived suppressor cells in breast cancer using single-cell transcriptomics. Sci Immunol 5, doi:10.1126/sciimmunol.aay6017 (2020).

25 Wu, N. et al. MerTK(+) macrophages promote melanoma progression and immunotherapy resistance through AhR-ALKAL1 activation. Sci Adv 10, eado8366, doi:10.1126/sciadv.ado8366 (2024).

26 Kim, D. et al. Ccr2+ Monocyte-Derived Macrophages Influence Trajectories of Acquired Therapy Resistance in Braf-Mutant Melanoma. Cancer Res 83, 2328–2344, doi:10.1158/0008-5472.Can-22-2841 (2023).

27 Vasilevska, J. et al. Monitoring melanoma patients on treatment reveals a distinct macrophage population driving targeted therapy resistance. Cell Rep Med 5, 101611, doi:10.1016/j.xcrm.2024.101611 (2024).

28 Mittal, S. K., Cho, K. J., Ishido, S. & Roche, P. A. Interleukin 10 (IL-10)-mediated Immunosuppression: MARCH-I INDUCTION REGULATES ANTIGEN PRESENTATION BY MACROPHAGES BUT NOT DENDRITIC CELLS. J Biol Chem 290, 27158–27167, doi:10.1074/jbc.M115.682708 (2015).

29 Sheban, F. et al. ZEB2 is a master switch controlling the tumor-associated macrophage program. Cancer Cell 43, 1227–1241.e1211, doi:10.1016/j.ccell.2025.03.021 (2025).

30 Balaban, R. et al. Efferocytosis induces proangiogenic function and chromatin remodeling in tumor-associated macrophages. Sci Immunol 11, eaed1544, doi:10.1126/sciimmunol.aed1544 (2026).

31 Yang, J. et al. Mature and migratory dendritic cells promote immune infiltration and response to anti-PD-1 checkpoint blockade in metastatic melanoma. Nat Commun 16, 8151, doi:10.1038/s41467-025-62878-5 (2025).

32 Suomalainen, A. & Nunnari, J. Mitochondria at the crossroads of health and disease. Cell 187, 2601–2627, doi:10.1016/j.cell.2024.04.037 (2024).

33 Lei, Y. et al. Cooperative sensing of mitochondrial DNA by ZBP1 and cGAS promotes cardiotoxicity. Cell 186, 3013–3032.e3022, doi:10.1016/j.cell.2023.05.039 (2023).

34 Holicek, P. et al. Type I interferon and cancer. Immunol Rev 321, 115–127, doi:10.1111/imr.13272 (2024).

35 Porritt, R. A. & Hertzog, P. J. Dynamic control of type I IFN signalling by an integrated network of negative regulators. Trends Immunol 36, 150–160, doi:10.1016/j.it.2015.02.002 (2015).

36 Falahat, R. et al. Epigenetic reprogramming of tumor cell-intrinsic STING function sculpts antigenicity and T cell recognition of melanoma. Proc Natl Acad Sci U S A 118, doi:10.1073/pnas.2013598118 (2021).

37 Wu, L. et al. KDM5 histone demethylases repress immune response via suppression of STING. PLoS Biol 16, e2006134, doi:10.1371/journal.pbio.2006134 (2018).

38 Suter, M. A. et al. cGAS–STING cytosolic DNA sensing pathway is suppressed by JAK2-STAT3 in tumor cells. Scientific Reports 11, 7243, doi:10.1038/s41598-021-86644-x (2021).

39 Scortegagna, M. et al. Age-Associated Modulation of TREM1/2-Expressing Macrophages Promotes Melanoma Progression and Metastasis. Cancer Res 85, 2218–2233, doi:10.1158/0008-5472.Can-24-4317 (2025).

40 Egan, D. et al. Chronic interferon signaling at baseline is associated with acquired resistance to immunotherapy in metastatic melanoma. Cell Rep Med 7, 102981, doi:10.1016/j.xcrm.2026.102981 (2026).

